# Asymmetric structures and conformational plasticity of the uncleaved full-length human immunodeficiency virus (HIV-1) envelope glycoprotein trimer

**DOI:** 10.1101/2021.03.28.437443

**Authors:** Shijian Zhang, Kunyu Wang, Wei Li Wang, Hanh T. Nguyen, Shuobing Chen, Maolin Lu, Eden P. Go, Haitao Ding, Robert T. Steinbock, Heather Desaire, John C. Kappes, Joseph Sodroski, Youdong Mao

**Affiliations:** Department of Cancer Immunology and Virology, Dana-Farber Cancer Institute, Boston, MA 02215, USA; Department of Microbiology, Harvard Medical School, Boston, MA 02115, USA; State Key Laboratory for Artificial Microstructures and Mesoscopic Physics, School of Physics, Center for Quantitative Biology, Peking University, Beijing 100871, China; Intel Parallel Computing Center for Structural Biology, Dana-Farber Cancer Institute, Boston, MA 02215, USA; Department of Microbial Pathogenesis, Yale University School of Medicine, New Haven, CT 06536, USA; Department of Chemistry, University of Kansas, Lawrence, KS 66049, USA; Department of Medicine, University of Alabama at Birmingham, AL 35294, USA; Birmingham Veterans Affairs Medical Center, Research Service, Birmingham, AL 35294, USA; Department of Immunology and Infectious Disease, Harvard T.H. Chan School of Public Health, Boston, MA 02115, USA

**Keywords:** Env, Cleavage, Furin, Processing, Conformation, cryo-electron microscopy, structure, antibody, asymmetry

## Abstract

The functional human immunodeficiency virus (HIV-1) envelope glycoprotein (Env) trimer [(gp120/gp41)_3_] is produced by cleavage of a conformationally flexible gp160 precursor. Gp160 cleavage or the binding of BMS-806, an entry inhibitor, stabilizes the pre-triggered, “closed” (State-1) conformation recognized by rarely elicited broadly neutralizing antibodies. Poorly neutralizing antibodies (pNAbs) elicited at high titers during natural infection recognize more “open” Env conformations (States 2 and 3) induced by binding the receptor, CD4. We found that BMS-806 treatment and crosslinking decreased the exposure of pNAb epitopes on cell-surface gp160; however, after detergent solubilization, crosslinked and BMS-806-treated gp160 sampled non-State-1 conformations that could be recognized by pNAbs. Cryo-electron microscopy of the purified BMS-806-bound gp160 revealed two hitherto unknown asymmetric trimer conformations, providing insights into the allosteric coupling between trimer opening and structural variation in the gp41 HR1_N_ region. The individual protomer structures in the asymmetric gp160 trimers resemble those of other genetically modified or antibody-bound cleaved HIV-1 Env trimers, which have been suggested to assume State-2-like conformations. Asymmetry of the uncleaved Env potentially exposes surfaces of the trimer to pNAbs. To evaluate the effect of stabilizing a State-1-like conformation of the membrane Env precursor, we treated cells expressing wild-type HIV-1 Env with BMS-806. BMS-806 treatment decreased both gp160 cleavage and the addition of complex glycans, implying that gp160 conformational flexibility contributes to the efficiency of these processes. Selective pressure to maintain flexibility in the precursor of functional Env allows the uncleaved Env to sample asymmetric conformations that potentially skew host antibody responses toward pNAbs.

**IMPORTANCE:** The envelope glycoprotein (Env) trimers on the surface of human immunodeficiency virus (HIV-1) mediate the entry of the virus into host cells and serve as targets for neutralizing antibodies. The functional Env trimer is produced by cleavage of the gp160 precursor in the infected cell. We found that the HIV-1 Env precursor is highly plastic, allowing it to assume different asymmetric shapes. This conformational plasticity is potentially important for Env cleavage and proper modification by sugars. Having a flexible, asymmetric Env precursor that can misdirect host antibody responses without compromising virus infectivity would be an advantage to a persistent virus like HIV-1.

## INTRODUCTION

Human immunodeficiency virus (HIV-1), the etiologic agent of acquired immunodeficiency syndrome (AIDS), utilizes a metastable envelope glycoprotein (Env) trimer to engage host receptors and enter target cells (1). The functional Env trimer consists of three gp120 exterior subunits and three gp41 transmembrane subunits (1–3). During virus entry, gp120 engages the receptors, CD4 and CCR5/CXCR4, and gp41 fuses the viral and cell membranes (4–16). Env is the only virus-specific protein on the viral surface and is targeted by host antibodies (17–20).

In infected cells, the HIV-1 Env trimer is synthesized in the rough endoplasmic reticulum (ER), where signal peptide cleavage, folding, trimerization and the addition of high-mannose glycans take place (21–24). The resulting gp160 Env precursor is transported to the Golgi apparatus, where some of the glycans are modified to complex types and proteolytic cleavage by host furin-like proteases produces the gp120 and gp41 subunits (25–41). The proteolytically processed, mature Env trimers are transported to the cell surface and incorporated into virions.

On the membrane of primary HIV-1, Env exists in a pre-triggered, “closed” conformation (State 1) that resists the binding of commonly elicited antibodies (42–47). Binding to the receptor, CD4, on the target cell releases the restraints that maintain Env in State 1, allowing transitions through a default intermediate conformation (State 2) to the pre-hairpin intermediate (State 3) (42,48,49). In the more “open” State-3 Env conformation, a trimeric coiled coil composed of the gp41 heptad repeat (HR1) region is formed and exposed, as is the gp120 binding site for the second receptor, either CCR5 or CXCR4 (50–58). Binding to these chemokine receptors is thought to promote the insertion of the hydrophobic gp41 fusion peptide into the target cell membrane and the formation of a highly stable six-helix bundle, which mediates viral-cell membrane fusion (14-16,59-62).

The ability of HIV-1 to establish persistent infections in humans requires an Env trimer that minimally elicits neutralizing antibodies and resists the binding of antibodies generated during the course of natural infection. In addition to a heavy glycan shield and surface variability, the conformational flexibility and plasticity of Env may help HIV-1 avoid the host antibody response (45,47,63–66). Flexible Envs could present epitopes that are not exposed on the State-1 Env trimer, misdirecting host antibodies away from the functional virus spike. The vast majority of antibodies elicited by Env during natural HIV-1 infection are unable to bind the functional State-1 Env trimer, and instead recognize downstream conformations (States 2, 2A and 3) (67–71). These antibodies cannot access their epitopes once the virus has bound CD4 and therefore do not neutralize efficiently (70). Uncleaved Envs that assume State-2/3 conformations are abundant on the surface of HIV-1-infected cells, in some cases reaching the cell surface by bypassing the Golgi (72). Poorly neutralizing antibodies (pNAbs) with State-2/3 specificity typically recognize these uncleaved Envs more efficiently than cleaved Env (73–79). Crosslinking the uncleaved cell-surface Env exerted effects on Env antigenicity similar to those resulting from gp120-gp41 cleavage, suggesting that the uncleaved Env might be more flexible than mature Env (80). Indeed, recent single-molecule fluorescence resonance energy transfer (smFRET) analysis confirmed that, in contrast to the dominant State-1 conformation of the wild-type Env, an Env mutant unable to be proteolytically processed due to an alteration of the cleavage site occupies States 2 and 3 more frequently than State 1 (81). Thus, the abundant, cell-surface-accessible and conformationally heterogeneous uncleaved Env could misdirect host immune responses away from the elicitation of broadly neutralizing antibodies, which generally recognize the State-1 Env conformation (42,45,46,48,81). Broadly neutralizing antibodies (bNAbs) typically appear after several years of HIV-1 infection and only in a minority of HIV-1-infected individuals (83–91).

Here, we investigate the conformation of the uncleaved HIV-1 Env trimer, both on the cell surface and purified from membranes. Cryo-electron microscopy (cryo-EM) reconstructions reveal that purified uncleaved Envs preferentially assume asymmetric trimer conformations, exposing epitopes for pNAbs. We identified a gp41 region in which structural changes are coupled to the asymmetric opening of the Env trimer. We tested the effect of a State-1-stabilizing gp120 ligand, BMS-378806 (herein called BMS-806) on the cleavage and glycosylation of the wild-type Env. Our findings indicate the importance of conformational plasticity of the uncleaved HIV-1 Env trimer for efficient proteolytic maturation, complex glycan addition and evasion of host antibody responses.

## RESULTS

### Analysis of the conformation of uncleaved HIV-1 Env on cell surfaces

Cleavage of the HIV-1 Env precursor affects its antigenicity (73–79). The recognition of the uncleaved and mature HIV-1_JR-FL_ Envs on the surface of transfected HOS cells exhibited distinct patterns for State 1-preferring bNAbs versus State 2/3-preferring pNAbs (Fig. 1A). Whereas the uncleaved Env was bound by antibodies capable of recognizing all three states, the mature Env was bound only by the potently neutralizing antibodies with State-1 preferences. The uncleaved Env apparently samples multiple conformations, but the mature Env assumes a conformation that precludes the binding of pNAbs.

**Figure 1.**
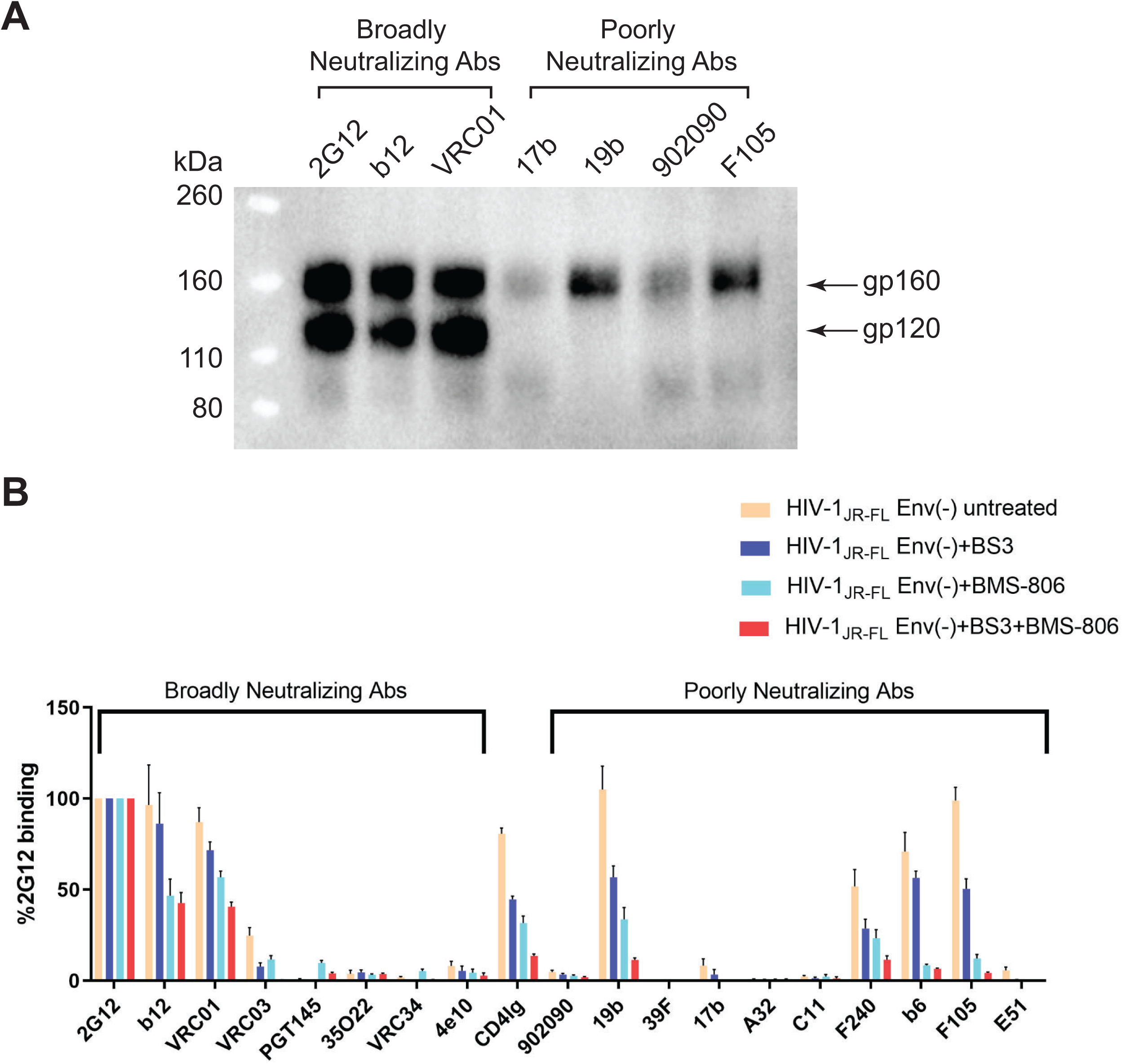
Antibody recognition of cleaved and uncleaved HIV-1 Envs on the cell surface. (A) HOS cells transiently expressing the wild-type HIV-1_JR-FL_ Env, a fraction of which is cleaved in these cells, were incubated with the indicated broadly neutralizing antibodies or poorly neutralizing antibodies. After washing and lysis of the cells, the antibody-Env complexes were purified with Protein A-Sepharose beads and analyzed by Western blotting with a rabbit anti-gp120 polyclonal serum. (B) The effect of crosslinking with BS3 and/or BMS-806 treatment on antibody binding to HIV-1_JR-FL_ Env(-) on the surface of CHO cells was evaluated by cell-based ELISA. BMS-806 (10 µM) was added to the CHO cells at the time of induction of Env(-) expression with doxycycline. All values were normalized against 2G12 antibody binding and were derived from at least three independent experiments. Note that the HIV-1_JR-FL_ Env(-) glycoprotein is not recognized by the PGT145 V2 quaternary antibody, which serves as a negative control.

The HIV-1 entry inhibitor, BMS-806, hinders transitions from State 1 and modestly increases the occupancy of State 1 by the mature, wild-type HIV-1 Env (see Table 1) (42,53,54,79,81). BMS-806 treatment or glutaraldehyde crosslinking has been shown to shift the antigenic profile of uncleaved HIV-1 Env closer to that of the cleaved Env (79, 80). Incubating virions containing uncleaved Env with BMS-806 significantly enriched the low-FRET State-1 conformation, resulting in a conformational profile closer to that of the unliganded mature HIV-1 Env (Table 1) (81). We tested the effects of BMS-806 and the lysine-specific crosslinker, bis (sulfosuccinimidyl) suberate (BS3), on the antigenic profile of cleavage-defective HIV-1_JR-FL_ Env(-) expressed on the surface of CHO cells (Fig. 1B). Treatment with BMS-806 and BS3 additively decreased Env(-) recognition by pNAbs (19b, b6, F105 and F240) and CD4-Ig, which preferentially bind Env conformations other than State 1 (45,48,54,78,81). In comparison, for the bNAbs 2G12, b12 and VRC01, the BMS-806/BS3-treated Env(-) was recognized at more than 40% the level observed for the untreated Env(-). These results are consistent with previous studies suggesting that BMS-806 can decrease the State-2/3 occupancy of uncleaved HIV-1 Envs anchored in the viral or cell membranes (Table 1) (79, 81).

**Table 1.** Summary of HIV-1_JR-FL_ conformational states^a^.

| Env | Source | Treatment | Occupancy of conformational states (%) |  |  | Reference |
| --- | --- | --- | --- | --- | --- | --- |
|  |  |  | State 1 | State 2 | State 3 |  |
| Wild-type HIV-1<sub>JR-FL</sub> Env | virion | None | 50 | 26 | 24 | 42,81 |
|  |  | BMS-806 | 55 | 18 | 27 |  |
| HIV-1<sub>JR-FL</sub> Env(-) | virion | None | 25 | 42 | 33 | 81 |
|  |  | BMS-806 | 40 | 32 | 28 |  |
|  | Purified from cell membranes | BMS-806 + BS3 crosslinking | 26 | 37 | 37 | This study |

<sup>a</sup>The relative occupancies (%) of conformational states for the indicated sources and treatments of HIV-1 Envs were derived from smFRET histograms. The FRET histograms were compiled from individual smFRET traces. The state distributions in the FRET histograms were fitted to the sum of three Gaussian distributions by hidden Markov modeling, and the occupancy of each state obtained from the area under each Gaussian curve.

### Purification and characterization of Env(-) trimers

To investigate further the range of conformations sampled by the uncleaved HIV-1 Env, we purified full-length HIV-1_JR-FL_ Env(-) trimers from the membranes of inducibly-expressing CHO cells (Fig. 2A and B). The CHO cells were incubated with BMS-806 during Env(-) synthesis in an attempt to shift occupancy from States 2/3 to State 1. BMS-806 treatment of the Env(-)-expressing cells reduced the synthesis of sialidase-sensitive and Endoglycosidase H-resistant glycoforms that are relatively enriched in complex carbohydrates (Fig. 2C). Glycosylation analysis revealed that BMS-806 treatment led to decreased complex sugar addition to the glycans modifying gp120 asparagine residues 88, 156, 160, 241, 362 and 463 (Fig. 2D and E). The effects of BMS-806 on Env(-) conformation apparently influence the conversion of particular high-mannose glycans to complex carbohydrates in the Golgi.

**Figure 2.**
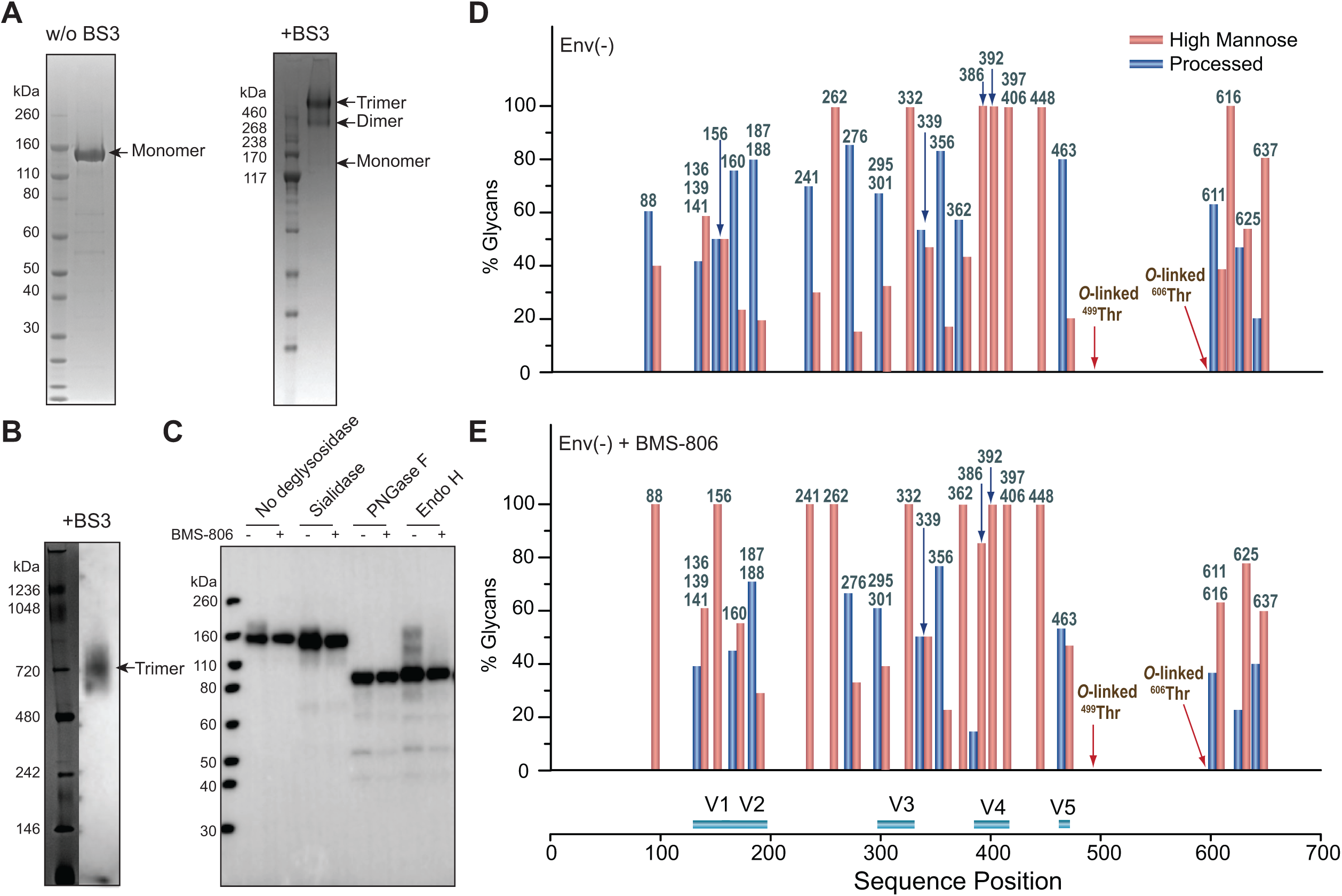
Characterization of the full-length HIV-1_JR-FL_ Env(-) glycoprotein in CHO cell lysates and in detergent-solubilized purified forms. (A) Purified HIV-1_JR-FL_ Env(-) without and with crosslinking by BS3 was run on a NUPAGE 4-12% BT gel stained by Coomassie Blue. (B) Purified HIV-1_JR-FL_ Env(-) crosslinked by BS3 was run on a NativePAGE 4-16%BT gel and subjected to Western blotting with an HRP-conjugated anti-HIV-1 gp120 antibody. (C-E) To evaluate the effect of BMS-806 on the glycosylation of the synthesized Env(-) glycoprotein, BMS-806 (10 µM) was added to the CHO cells at the time of doxycycline induction. (C) The effect of BMS-806 on HIV-1_JR-FL_ Env(-) glycosylation was evaluated by Western blotting after digestion with glycosidases (sialidase, Peptide-N-glycosidase F (PNGase F), and Endoglycosidase H (Endo H)). The purified HIV-1_JR-FL_ Env(-) glycoproteins were digested with the indicated glycosidase, run on a NUPAGE 4-12% BT gel, and subjected to Western blotting with an HRP-conjugated anti-HIV-1 gp120 antibody. The results shown are representative of those obtained in three independent experiments. Note that BMS-806 treatment decreases Env(-) heterogeneity by reducing the levels of sialidase-sensitive and Endo H-resistant glycoforms. (D,E) The bar graphs show the glycan profiles at each glycosylation site of HIV-1_JR-FL_ Env(-) purified from untreated CHO cells (D) or CHO cells treated with 10 µM BMS-806 (E), as determined by mass spectrometry. The glycan composition (in percent) was broadly characterized as high-mannose (red bars) or processed (complex + hybrid) glycans (blue bars).

To purify the Env(-) trimer complexes, membranes from BMS-806-treated CHO cells were incubated with saturating concentrations of BMS-806, crosslinked with BS3, and solubilized in Cymal-5. The detergent in the Env(-) glycoprotein solution was exchanged to a mixture of 4.5 mg/ml amphipol A8-35 and 0.005% Cymal-6 prior to cryo-plunging the samples in preparation for eventual cryo-electron microscopy (cryo-EM) imaging. Parallel smFRET studies estimated that only 26% of detergent-solubilized Env(-) was in a low-FRET conformation consistent with State 1 (Fig. 3A). The majority (74%) of the solubilized Env(-) glycoproteins assumed high- and intermediate-FRET conformations consistent with States 2 and 3, respectively. Thus, compared with BMS-806-treated Env(-) on virions, the Env(-) glycoproteins solubilized and purified from CHO cells exhibit less State 1 and more State 2/3 conformations (Table 1).

**Figure 3.**
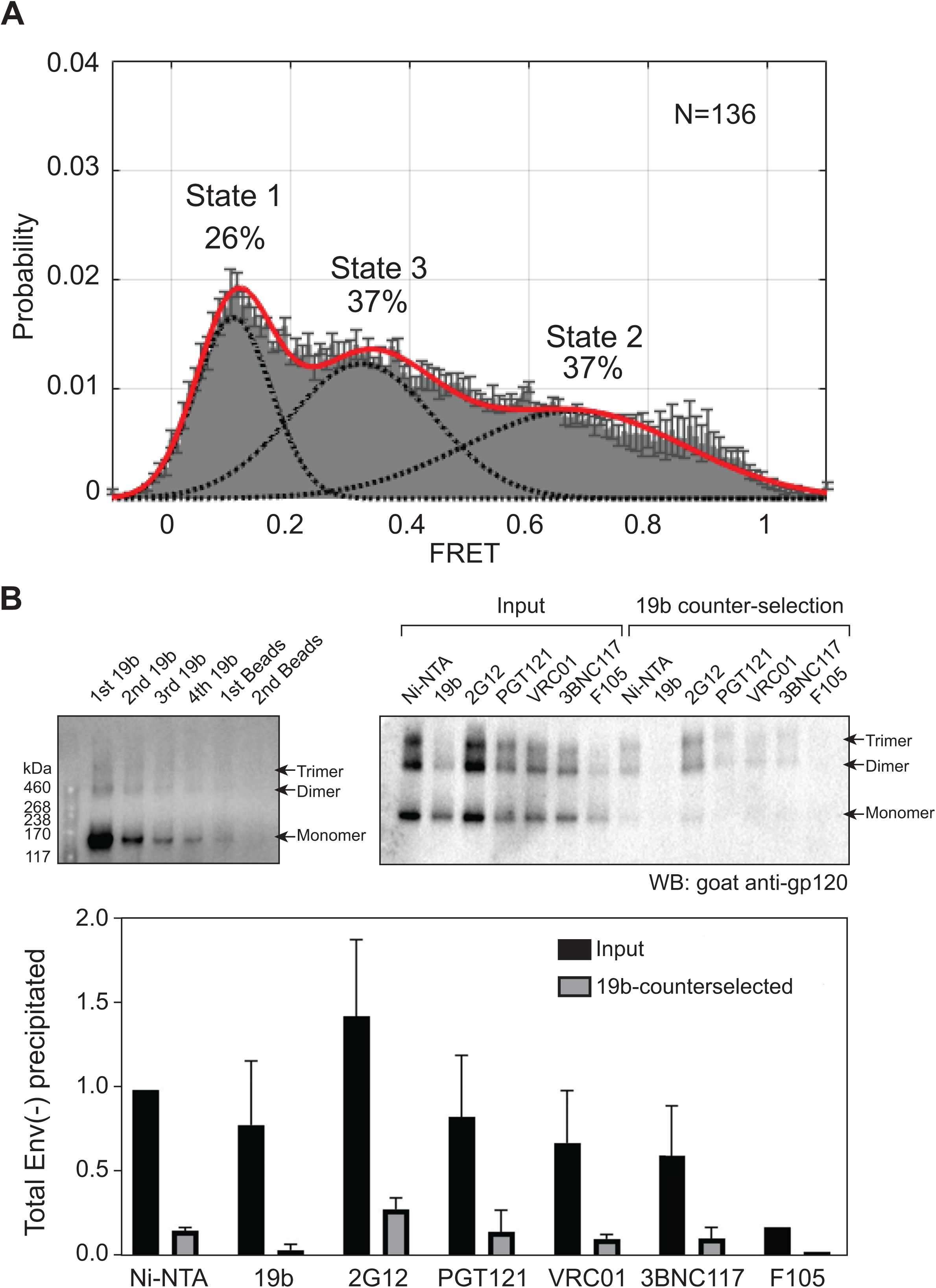
Conformations of purified HIV-1_JR-FL_ Env(-) treated with BMS-806 and crosslinked with BS3. (A) HIV-1_JR-FL_ Env(-) with V1 and V4 labeling tags was purified from 293T cell membranes using a protocol identical to that used for preparation of Env(-) for cryo-EM imaging. The purified Env(-) was labeled and analyzed by smFRET. FRET trajectories were compiled into a population FRET histogram and fit to the Gaussian distributions associated with each conformational state, according to a hidden Markov model (42). The percentage of the population that occupies each state as well as the number of molecules analyzed (N) is shown. The error bars represent the standard deviation from three independent data sets. (B) Membranes from BMS-806-treated CHO cells expressing HIV-1_JR-FL_ Env(-) were crosslinked with BS3 and then solubilized in Cymal-5 detergent. The lysate was successively incubated with the 19b anti-gp120 (V3) antibody and Protein-A Sepharose beads. The Env(-) glycoproteins precipitated by the 19b antibody or by the negative-control Protein A-Sepharose beads during the indicated rounds of immunoprecipitation were analyzed by SDS-PAGE and Western blotting (upper left panel). The Env(-) glycoproteins in the initial cell membrane lysate (Input) and those glycoproteins remaining after four rounds of 19b counterselection were precipitated with Ni-NTA beads or the indicated antibodies; the precipitated Env(-) glycoproteins were analyzed by SDS-PAGE and Western blotting (upper right panel). The total amounts of Env(-) glycoprotein in the input and after 19b counterselection, normalized to the input Env(-) glycoprotein amount precipitated by the Ni-NTA beads, are shown in the bar graph (lower panel). Means and standard deviations derived from two independent experiments are shown.

The increased exposure of the gp120 V3 loop is a sensitive indicator of HIV-1 Envs that have undergone transitions from a State-1 conformation (48,54,92–94). We tested the ability of the 19b anti-V3 antibody, which does not neutralize most primary HIV-1 strains, to precipitate the BMS-806-treated, BS3-crosslinked Env(-) trimers solubilized in Cymal-5 detergent (Fig. 3B). After successive precipitations with the 19b antibody, approximately 85% of the Env(-) glycoprotein was removed from the CHO cell lysate. Therefore, even in the presence of BMS-806 and after BS3 crosslinking, most of the Env(-) trimers solubilized in Cymal-5 detergent apparently sample non-State-1 conformations. Together with the above cell-based ELISA and smFRET results, these experiments suggest that detergent solubilization destabilizes the uncleaved Env, even after BMS-806 and BS3 treatment. Therefore, the cell membrane and lipid-protein interactions may be important for the stabilization of the Env(-) State-1 conformation.

### Env(-) structure determination by cryo-electron microscopy (cryo-EM)

The BMS-806-treated, BS3-crosslinked HIV-1_JR-FL_ Env(-) trimers, purified in Cymal-5 and exchanged into amphipol A8-35 and Cymal-6, were analyzed by cryo-EM. We collected cryo-EM data from both a 200-kV FEI Tecnai Arctica microscope without an energy filter and a 300-kV FEI Titan Krios microscope with a Gatan BioQuantum energy filter, in video frames of a super-resolution counting mode with the Gatan K2 Summit direct electron detector (Fig. 4A-F). While both 200-kV and 300-kV cryo-EM datasets gave rise to consistent results, the final reconstructions at near-atomic resolution were achieved using the 300-kV cryo-EM dataset; the 300-kV dataset incorporates single-particle data collected at a high tilt angle of the sample stage to alleviate the effect of the strong orientation preference of the Env(-) particles. By contrast, the 200-kV cryo-EM dataset, which lacks tilted data, fell short of achieving a comparable level of resolution and suffered from the orientation preference of the particle images. However, despite the modest level of resolution (5.5-8 Å), extensive 3D classification of the 200-kV dataset, as detailed in a bioRxiv preprint (95), indicated the existence of multiple Env(-) conformations, some of which are consistent with the higher-resolution reconstructions obtained from the 300-kV dataset. This paper focuses on interpreting two higher-resolution maps of the uncleaved Env(-) trimer derived from the 300-kV dataset.

**Figure 4.**
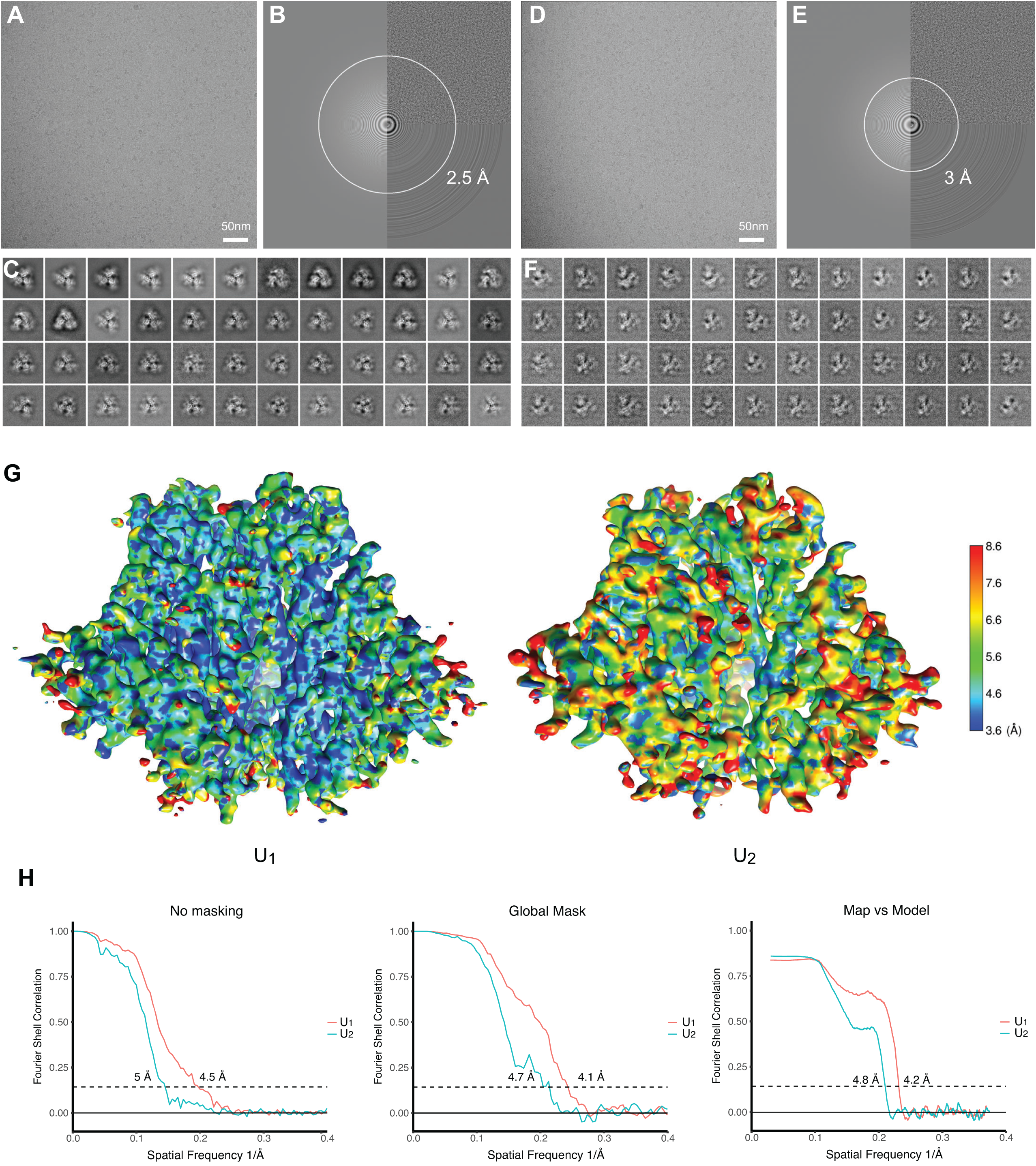
Cryo-EM analysis of the full-length HIV-1_JR-FL_ Env(-) trimer. (A) A typical cryo-EM micrograph of Env(-) trimers taken with a Gatan K2 direct electron detector at 0 degrees of tilt. (B) Fourier transform of the image in A . (C) Unsupervised 2D class averages for non-tilt particles. (D) A typical cryo-EM micrograph of Env(-) trimers taken with a Gatan K2 direct electron detector at 45 degrees of tilt. (E) Fourier transform of the image in D. (F) Unsupervised 2D class averages for tilted particles. (G) The local resolution measurement of the State-U_1_ and State-U_2_ maps, as measured by ResMap (143). The maps are colored according to the local resolution, indicated by the color gradient (units in Angstroms). Side views of the Env(-) maps are shown, with gp120 at the bottom of the figure and gp41 at the top. (H) The gold-standard FSC plots of the State-U_1_ and State-U_2_ cryo-EM maps.

Analysis of the 300-kV data resulted in two major 3D classes, herein designated State U_1_ and State U_2_, respectively comprising 37% and 17% of the imaged particles, after removal of junk particles. The State-U_1_ and State-U_2_ maps were refined to 4.1 and 4.7 Å, respectively, without imposing any symmetry during refinement and reconstruction (Fig. 4G and H). The map quality allowed atomic modelling and refinement with accuracy to the level of the Cα backbone trace. By contrast, imposing C3 symmetry during refinement and reconstruction resulted in lower resolution and poorer structural features in the refined density maps of both State-U_1_ and State-U_2_, suggesting that both conformations indeed lack rigorous three-fold symmetry. Other 3D classes derived from the HIV-1_JR-FL_ dataset were not able to be refined to comparable levels of resolution, and thus are not further analyzed and discussed herein. Curiously, no major 3D classes with rigorous three-fold symmetry were found when extensive 3D classification was conducted using the maximum-likelihood method without imposing C3 symmetry (96). This likely reflects the intrinsic conformational plasticity of the Env(-) glycoprotein, although we do not rule out the contribution of preparation-dependent variables, such as asymmetric crosslinking between adjacent protomers.

### Key structural features of the asymmetric uncleaved HIV-1 Env trimers

The U_1_ and U_2_ Env(-) trimers share an overall topology with existing structures of soluble and membrane HIV-1 Env trimers (97–108) (Fig. 5A). A central feature of all these structures is a 3-helix bundle (3-HB_C_) formed by the C-terminal portion of the gp41 HR1 region (HR1_C_); the gp120 subunits project outward from this central helical coiled coil. These common features allowed us to use existing symmetric and asymmetric HIV-1 Env trimer structures as references to build structural models of states U_1_ and U_2_. All three individual protomers in the U_1_ and U_2_ trimers exhibit similar folds, with Cα RMSD values of ∼2 Å (Fig. 5B and C). While both the U_1_ and U_2_ conformations of Env(-) are asymmetric, they exhibit different degrees of such asymmetry in terms of the relative rotation of the individual protomers with respect to the trimer axis. The protomers are differentially translated and rotated with respect to each other in unique ways in the U_1_ and U_2_ trimers (Fig. 5D), generating ∼3-4 Å movement overall in the gp120 outer domain (OD). When one of the protomers is used to align both conformations, the other two protomers of U_2_ are notably rotated by 2.8 and 4 degrees relative to the corresponding protomers of U_1_ (Fig. 5A). This creates the smallest and largest openings between two adjacent protomers in U_2_, the more asymmetric of the two Env(-) conformations. Alignment of all three protomer structures in each conformation indicates that the asymmetric conformations are facilitated by local structural rearrangements of residues 546-568 at the inter-protomer interface. This gp41 segment (HR1_N_) is immediately N-terminal to the central 3-HB_C_ and exhibits the greatest local structural variation among the promoters. Notably, the overall structural variation of gp41 among the U_1_ and U_2_ protomers is greater than that of the gp120 core structure, presumably because gp41 contributes more interactions to the inter-protomer interface. Consistently, the gp120 trimer association domain (TAD), which includes the V1/V2 and V3 regions, exhibits greater conformational variation in U_2_ than in U_1_, leading to an overall greater extent of asymmetry in U_2_ (Fig. 5B and C). There is similarly greater gp41 structural variation among the protomers in U_2_ than in U_1_.

**Figure 5.**
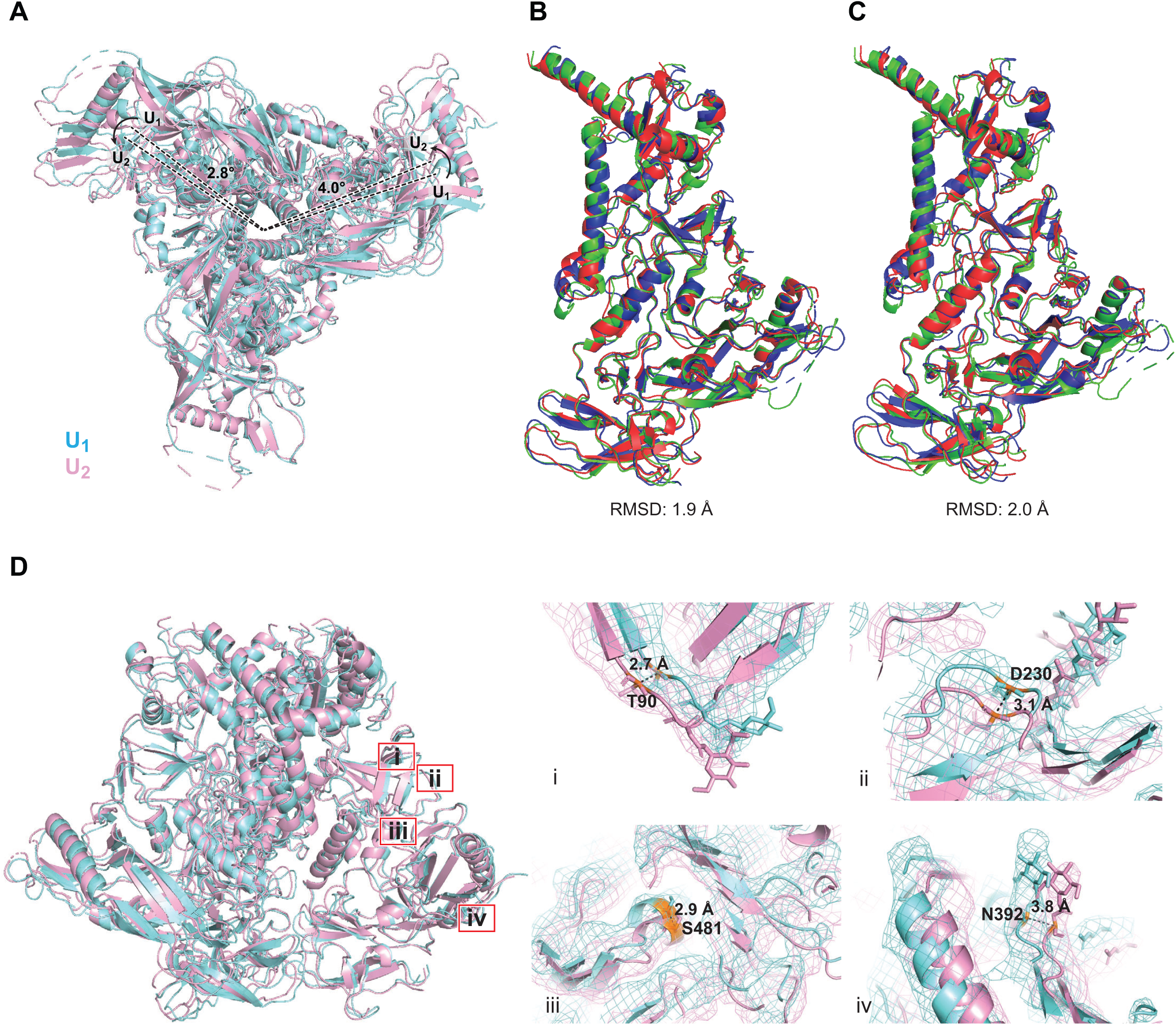
Comparison of U_1_ and U_2_ Env(-) structures. (A) Protomer 2 of the State-U_1_ and State-U_2_ models are superposed, showing that protomer 1 and protomer 3 are rotated 4.0° and 2.8°, respectively. (B) Three protomers of the State-U_1_ model are superposed. (C) Three protomers of the State-U_2_ model are superposed. (D) With protomer 2 of the State-U_1_ and State-U_2_ models superposed, the Cα distances between the same residues on the U_1_ and U_2_ structures are measured for four residues (from (i) to (iv): T90, D230, S481 and N392). In the side views of Env(-) shown in B-D, gp120 is at the bottom of the figure and gp41 at the top.

### Comparison with structures of cleaved HIV-1 Env trimers

We compared the U_1_ and U_2_ HIV-1_JR-FL_ Env(-) structures to those of mature (cleaved) HIV-1 Env trimers. The structure of the unliganded HIV-1_BG505_ sgp140 SOSIP.664 glycoprotein (PDB: 4ZMJ) provides an example of a stabilized soluble Env trimer with C3 symmetry (104). Structures of cytoplasmic tail-deleted, detergent-solubilized HIV-1_JR-FL_ and HIV-1_AMC011_ EnvΔCT trimers have been solved in complex with Fab fragments of the PGT151 neutralizing antibody (PDB: 5FUU and 6OLP, respectively) (105, 108). Binding of the PGT151 Fabs introduces asymmetry into the Env trimer, limiting the binding stoichiometry to two Fabs per trimer.

The folds of the U_1_ and U_2_ Env(-) protomers resemble those of the sgp140 SOSIP.664 and PGT151-bound EnvΔCT protomers (Fig. 6). The largest structural difference is localized in HR1_N_ residues 534-570 leading to the central 3-HB_C_ of gp41. When the U_1_ and sgp140 SOSIP.664 trimer structures are aligned using one of the protomers, the other two protomers of U_1_ exhibit rotations in opposite directions relative to the symmetric sgp140 SOSIP.664 trimer structure, causing a prominent narrowing of the opening angle between these two protomers in the U_1_ trimer structure (Fig. 6A). By contrast, when the U_1_ structure is aligned to the PGT151-bound EnvΔCT trimer using the protomer free of the antibody, both the other two protomers exhibit rotations in the same direction; this results in two smaller opening angles and one notably larger opening angle in comparison with those seen in the symmetric sgp140 trimer (Fig. 6B). In addition to relative rotation, the gp120 components of the U_1_ protomers also exhibit outward movement in both comparisons (Fig. 6A and B), giving rise to a slightly wider trimer footprint (Fig. 7A). Some local divergence in the gp120 V1/V2 region and gp41 α8 helix between HIV-1_JR-FL_ Env(-) and HIV-1_BG505_ sgp140 SOSIP.664 likely results from strain-dependent differences in primary sequence. Consistent with this explanation, the protomer structures of the Env(-) and EnvΔCT trimers, both derived from the HIV-1_JR-FL_ strain, align well in these regions. As is the case for all current HIV-1 Env trimer structures, the gp41 membrane-proximal external region (MPER) and transmembrane region are disordered in the U_1_ and U_2_ maps.

**Figure 6.**
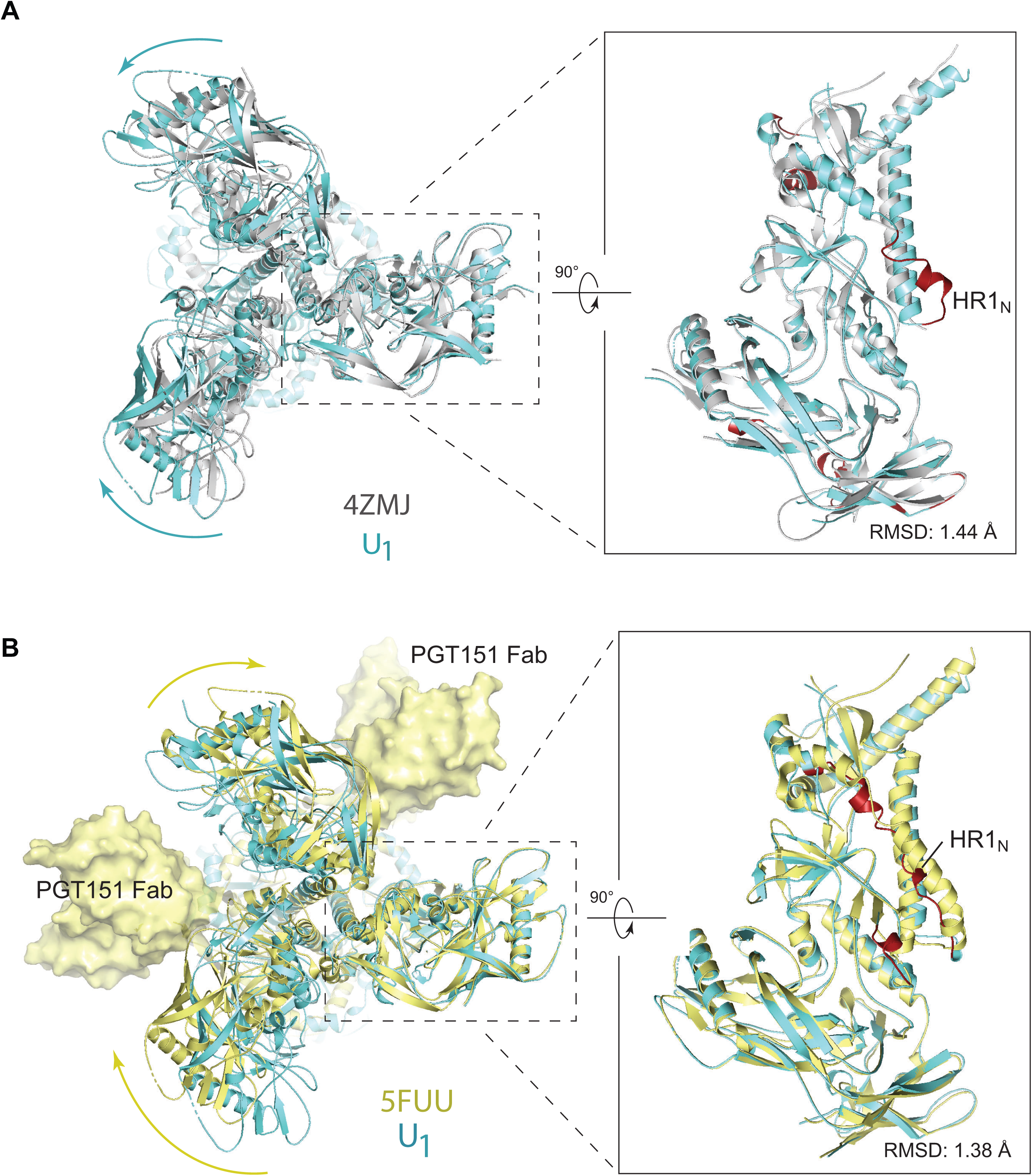
Comparison of Env(-) structures with those of cleaved HIV-1 Envs. (A) Left: Protomer 1 of the State-U_1_ trimer is superposed on the unliganded HIV-1_BG505_ sgp140 SOSIP.664 trimer (PDB ID 4ZMJ) (104), demonstrating how the other two protomers in State-U_1_ are rotated towards each other. Right: Side views of the superposed protomers, with red parts representing the major areas of difference between the two protomers. (B) Left: Protomer 1 of the State-U_1_ trimer is superposed on the HIV-1_JR-FL_ EnvΔCT trimer complexed with PGT151 Fabs (PDB ID 5FUU) (105), indicating that binding of the PGT151 antibodies introduces asymmetry into the Env trimer that differs from that of U_1_. Right: Side views of the superposed protomers, with red parts representing the major areas of difference between the two protomers. In the side views of the Env protomers shown in the right-hand panels of A and B, gp120 is at the bottom of the figure and gp41 at the top.

**Figure 7.**
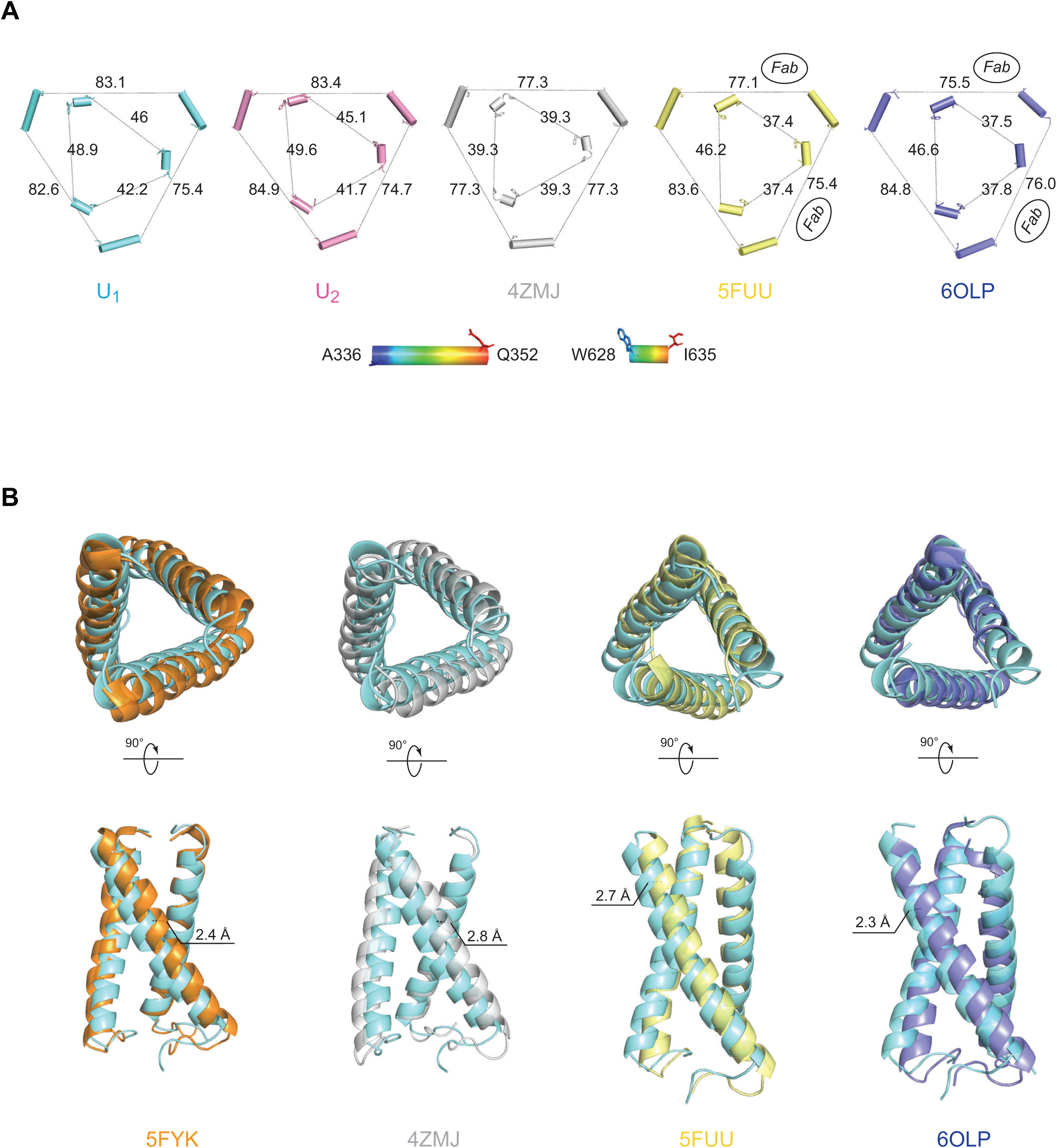
Comparison of Env trimer geometry among Env(-) trimers and mature Env trimers. (A) The inter-protomer distances (in Å) between selected atoms of gp120 and gp41 are shown for different Env structures: the smaller, inner triangle depicts distances measured between gp41 residues W628 and I635; the larger, outer triangle depicts distances measured between gp120 residues A336 and Q352. The U_1_ and U_2_ structures are compared with those of the unliganded sgp140 SOSIP.664 trimer (PDB ID 4ZMJ) (104) and the PGT151-bound HIV-1_JR-FL_ and HIV-1_AMC011_ EnvΔCT trimers (PDB IDs 5FUU and 6OLP, respectively) (105,108). For 5FUU and 6OLP, the sides of the Env trimer that are bound by the PGT151 Fabs are marked. (B) The three gp120 subunits of four Env trimer atomic structures were superposed with the gp120 subunits of the State-U_1_ Env(-) trimer. Each protomer was aligned separately. After gp120 alignment, the relative positions of the gp41 HR1_C_ helixes are jointly shown here. In each case, the U_1_ HR1_C_ helices are colored cyan. With gp120 aligned, the gp41 in State U_1_ is displaced compared with the other structures. Upper row: top views of 3-helix bundles; Bottom row: side views of 3-helix bundles. 5FYK is the structure of an HIV-1_JR-FL_ sgp140 SOSIP.664 trimer complexed with several neutralizing antibody Fabs (65).

We next compared the topology of the Env(-) trimers to that of cleaved Env trimers. The inter-protomer distances between arbitrarily chosen atoms on the outer surface of gp120 and gp41 provide a measure of trimer geometry (Fig. 7A). Of the trimers that we compared, the symmetric HIV-1_BG505_ sgp140 SOSIP.664 trimer is the most tightly packed, with the respective gp120 and gp41 sides 77.3 and 39.3 Å in length. The two sides of the EnvΔCT trimers bound to the PGT151 antibody Fabs are similar in length (gp120: 75.4, 77.1/gp41: 37.4, 37.4 Å and gp120: 75.5, 76.0/gp41: 37.5, 37.8 Å in the HIV-1_JR-FL_ and HIV-1_AMC011_ EnvΔCT trimers, respectively); these Fab-bound sides are shorter than the “opened” unliganded side (gp120: 83.6/gp41: 46.2 Å and gp120: 84.8/gp41: 46.6 Å in the HIV-1_JR-FL_ and HIV-1_AMC011_ EnvΔCT trimers, respectively). The asymmetry of the U_1_ Env(-) trimer is qualitatively similar to that of the U_2_ trimer; the asymmetry of the Env(-) trimers is distinguished by three sides of different lengths and therefore differs from the asymmetry in the EnvΔCT trimers induced by the PGT151 antibody. Notably, the average lengths of the gp120/gp41 sides of the Env(-) trimers are longer than those of the unliganded sgp140 SOSIP.664 or PGT151-bound EnvΔCT trimers, indicating that the uncleaved Env(-) trimers are packed less tightly than the cleaved Env trimers.

**To evaluate the basis for the increased “openness” of the uncleaved Env**(-) **trimers, we compared the structures of the gp41 3-HB_C_ coiled coil and HR1_N_ region in the Env**(**-**) **and cleaved Env trimers. Changes in the packing or orientation of the 3-HB_C_ coiled coil could potentially influence trimer topology. Although it appears that the crossing angles between two adjacent helices in the gp41 3-HB_C_ coiled coil are very similar in the U_1_ and U_2_ trimers, these 3-HB_C_ helices exhibit differential packing and asymmetric features in U_1_ and U_2_ that are amplified into a greater degree of overall trimeric asymmetry. Compared to the PGT151-bound cleaved Env structures (PDB IDs 5FUU and 6OLP), the U_1_ conformation has clearly larger crossing angles and thus a greater 3-HB_C_ coiled-coil footprint (Fig. 7B). By contrast, the crossing angles in U_1_ are nearly identical to those of the sgp140 SOSIP.664 trimers, but the U_1_ 3-HB_C_ helices exhibit marked translation in opposite directions that breaks the trimer symmetry seen in the crystal structures of the sgp140 SOSIP.664 trimers (PDB IDs 5FYK and 4ZMJ). Being able to sustain structural rearrangements involving both of the orthogonal degrees of freedom demonstrates that the Env trimer metastability and lability is potentially rooted in the conformational plasticity and flexibility of the central 3-HB_C_ structure.**

Despite a high degree of primary sequence conservation among HIV-1 strains, the gp41 HR1_N_ region (residues 541-570) exhibits significant conformational polymorphism among current HIV-1 Env trimer structures. In the pre-triggered (State-1) Env conformation, the gp41 HR1_N_ region has been implicated in the non-covalent association with gp120; in the pre-hairpin intermediate (State 3), the HR1_N_ region forms part of the extended HR1 helical coiled coil (14-16,109-111). Therefore, HR1_N_ may transition from an as-yet-unknown State-1 structure to a helical coiled coil (State 3) as Env “opens” upon binding CD4. The HR1_N_ region is relatively disordered in most sgp140 SOSIP.664 structures, probably as a result of the I559P change used to stabilize these soluble Env trimers (112–115). Even in asymmetric structures of sgp140 SOSIP.664 trimers bound to soluble CD4 and the E51 CD4-induced antibody (116), HR1_N_ disorder precludes analysis. We therefore limited our comparison to asymmetric Env trimers for which HR1_N_ structural information is available. Comparison of the HR1_N_ conformation in the asymmetric Env trimers suggested that the helicity of the HR1_N_ region is related to the degree of “openness” of the corresponding protomer (Fig. 8). **Lower helicity of the HR1_N_ region leads to a somewhat collapsed conformation that is correlated with a smaller inter-protomer opening angle. This is consistent with the notion that a non-helical, loop-like and more collapsed HR1_N_, which is located in the crevice formed by the protomer arms, would not have sufficient structural strength to sustain a wider opening angle. These observations support the proposition that the HR1_N_ conformation is allosterically coupled with asymmetric features of the 3-HB_C_ and the overall asymmetry of the entire trimer.**

**Figure 8.**
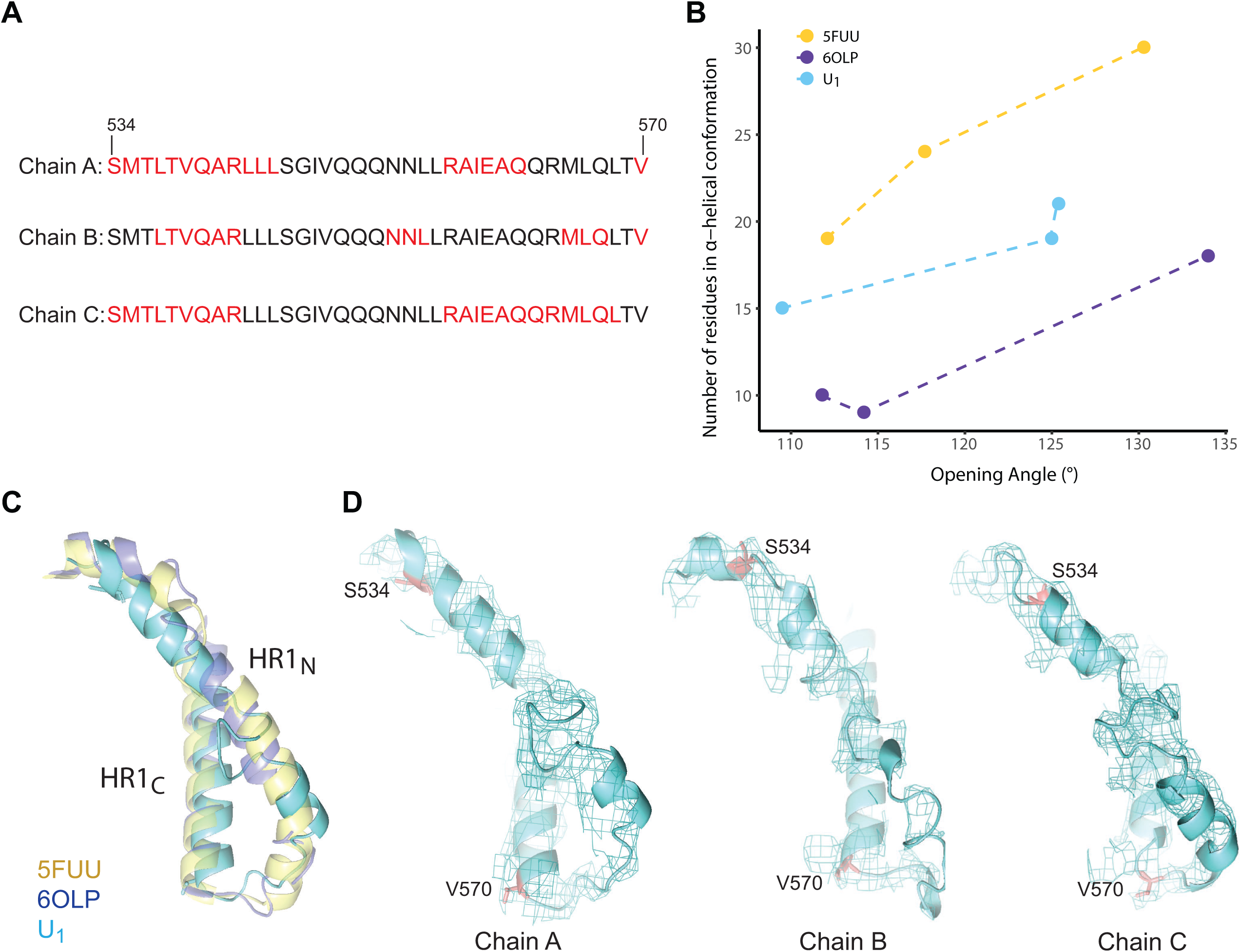
Relationship between HR1_N_ helicity and the opening angle of the trimer. (A) Sequences of the gp41 HR1_N_ region from three U_1_ protomers are shown, with residues in α-helices highlighted in red. (B) The relationship between HR1_N_ helicity and the opening angle of three asymmetric HIV-1 Env trimers (U_1_ and two PGT151-Fab-bound EnvΔCT trimers (PDB IDs 5FUU and 6OLP)) is shown. The x-axis represents the opening angle for each of three sides, measured using the “angle_between_domains” command in Pymol (142). The y-axis represents the number of residues in an α-helical conformation for the HR1_N_ region of that side. (C) The HR1_N_ and HR1_C_ regions from the three indicated atomic models are superposed. (D) The HR1_N_ regions from the three protomers in State U_1_ are shown .

### Env(-) glycosylation

Most of the peptide-proximal density associated with N-linked glycosylation is preserved in the U_1_ map and was modeled (Fig. 9). Most distal glycan residues are not well resolved, reflecting their dynamic nature and heterogeneity. As has been previously shown, the high-mannose glycans are clustered in a patch on the surface of the gp120 outer domain (39,40,65,117). No glycan-associated density on Asn 297 is detectable, and the glycan signal on Asn 448 is weak. The signals associated with the complex glycans on gp41 residues Asn 611 and Asn 637 are buried in noise. The most membrane-proximal gp41 glycan on Asn 616 is largely modified by high-mannose glycans.

**Figure 9.**
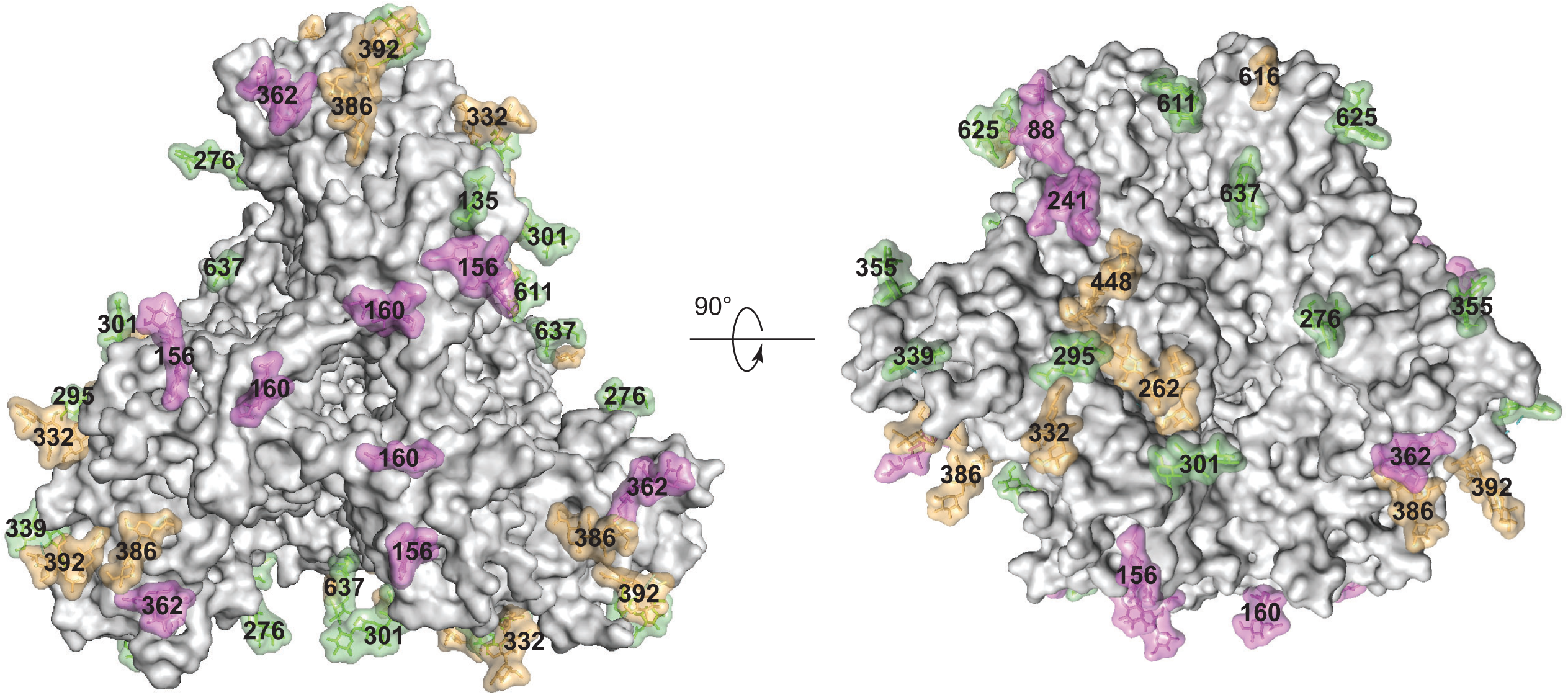
HIV-1_JR-FL_ Env(-) glycan structure. Glycans on State-U_1_ trimers are colored according to the following scheme: glycans that exhibit significant decreases in the addition of processed glycans as a result of BMS-806 treatment are colored purple; high-mannose glycans are colored yellow; and the remaining mixed or processed glycans that are not affected by BMS-806 binding are colored green.

BMS-806 treatment of Env(-)-expressing cells led to a reduction in the modification of glycans on Asn 88, 156, 160, 241, 362 and 463. Asn 88 and 241 are located at the gp120-gp41 interface, and Asn 156 and 160 at the trimer apex (Fig. 9). Previous studies have suggested that BMS-806 can strengthen inter-subunit and inter-protomer interactions in the Env trimer, increasing the binding of neutralizing antibodies that recognize the gp120-gp41 interface and trimer apex (79). Strengthening these interactions may render the carbohydrates in these regions less available for modification to complex carbohydrates. Consistent with this, two other BMS-806-sensitive glycans (on Asn 362 and Asn 463) reside on the perimeter of the gp120 outer domain that, in a more closed trimer, might be sterically limited by inter-protomer effects.

### BMS-806 binding site

The binding site of BMS-806 in sgp140 SOSIP.664 complexes (PDB: 5U7M) has been previously characterized (118). In the Env(-) maps, density corresponding to the location of BMS-806 in the sgp140 SOSIP.664 complexes is evident. In the Env(-) complexes, BMS-806 is located in the gp120 Phe 43 cavity and the adjacent water-filled channel, sandwiched between Trp 427 and Trp 112. Although the level of resolution does not allow unambiguous definition of the binding mode, the position and orientation of BMS-806 is consistent with that in the sgp140 SOSIP.664 complexes (118) (Fig. 10). In the U_1_ Env(-)-BMS-806 structure, as in the unliganded and BMS-806-bound sgp140 SOSIP.664 structures (104, 118), Layer 1 of the gp120 inner domain appears to be stabilized by the insertion of Trp 69 into the back end of the Phe 43 cavity, where it interacts orthogonally with Trp 112. During the course of Env binding to CD4, Layer 1 is thought to undergo rearrangement to decrease the off-rate of CD4 (119); fixation of Layer 1 by BMS-806 could help to inhibit Env conformational transitions to the CD4-bound State 3.

**Figure 10.**
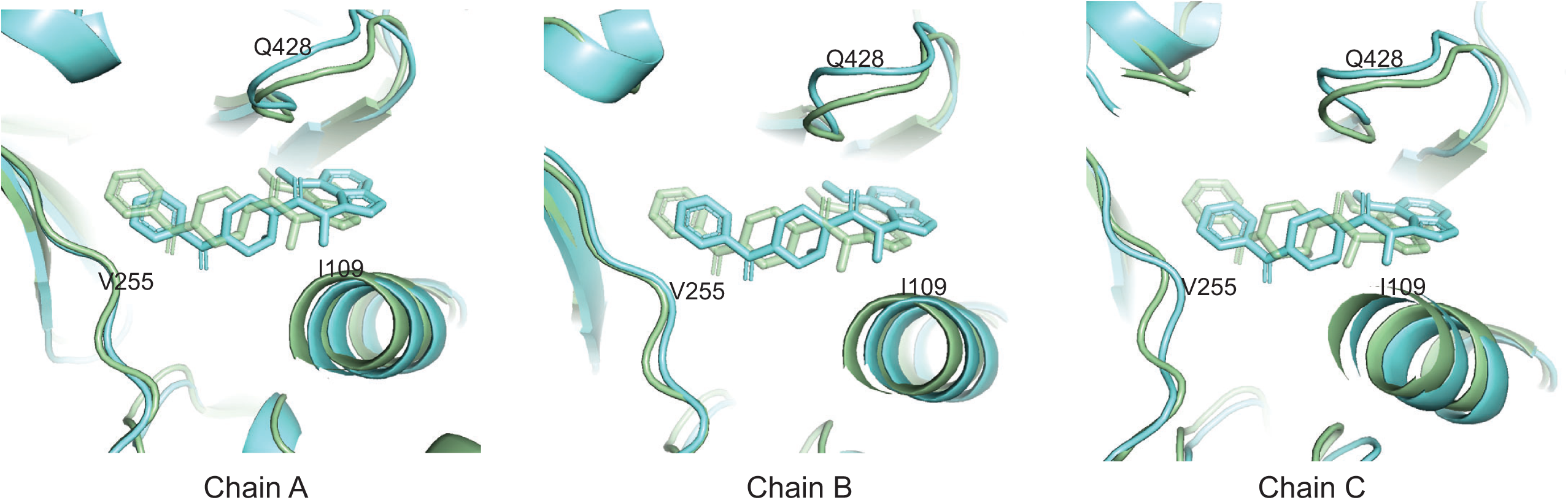
BMS-806 binding site. The BMS-806 binding sites within three protomers of the State-U_1_ structure (cyan) are compared with those in the BMS-806-bound sgp140 SOSIP.664 trimer (PDB 5U70) (118).

### Effect of BMS-806 on processing of wild-type HIV-1 Env

BMS-806 and its analogues block transitions from the pre-triggered Env conformation; thus, addition of these compounds to cleaved and uncleaved Envs on virions enriches State 1 (Table 1) (42,53,54,79,81). The studies shown in Figure 2D and E suggest that limiting the conformational flexibility of the cleavage-defective Env(-) by exposing Env(-)-expressing cells to BMS-806 can influence the processing of carbohydrate structures. To evaluate more thoroughly how Env conformation influences its processing, we used A549-Gag/Env cells, which produce virus-like particles (VLPs) containing Env (72). The wild-type HIV-1_AD8_ Env in the A549-Gag/Env cells is proteolytically processed and the VLPs contain mostly cleaved Env, as is the case for authentic HIV-1 virions (72). Therefore, the use of A549-Gag/Env cells allowed us to evaluate the effects of BMS-806 on the cleavage and glycosylation of wild-type HIV-1 Env in cells and on VLPs.

We incubated A549-Gag/Env cells with BMS-806 and studied Env in cell lysates and VLPs. BMS-806 treatment during Env expression resulted in a decrease in the efficiency of Env cleavage (Fig. 11A). The uncleaved Env produced in the presence of BMS-806 was efficiently incorporated into VLPs (Fig. 11B). This contrasts with the relative exclusion of uncleaved Env from VLPs produced in the absence of BMS-806 (Fig. 11B) (72). In the untreated cells, some of the glycans on the uncleaved Env are Endoglycosidase Hf-resistant and therefore are complex carbohydrates (Fig. 11A). The Endoglycosidase Hf-resistant fraction of the uncleaved Env migrated faster on SDS-polyacrylamide gels following BMS-806 treatment, indicating that fewer complex sugars are added to Env produced in A549-Gag/Env cells treated with BMS-806 (Fig. 11A). Nonetheless, in the BMS-806-treated cells, the uncleaved Env that is modified by complex glycans (and therefore has passed through the Golgi) is incorporated into VLPs (Fig. 11B). These results suggest that the BMS-806-induced reduction in the conformational flexibility of the Env precursor decreases the efficiency of gp160 cleavage and addition of some complex glycans, without significantly affecting Env transport through the Golgi or incorporation into VLPs.

**Figure 11.**
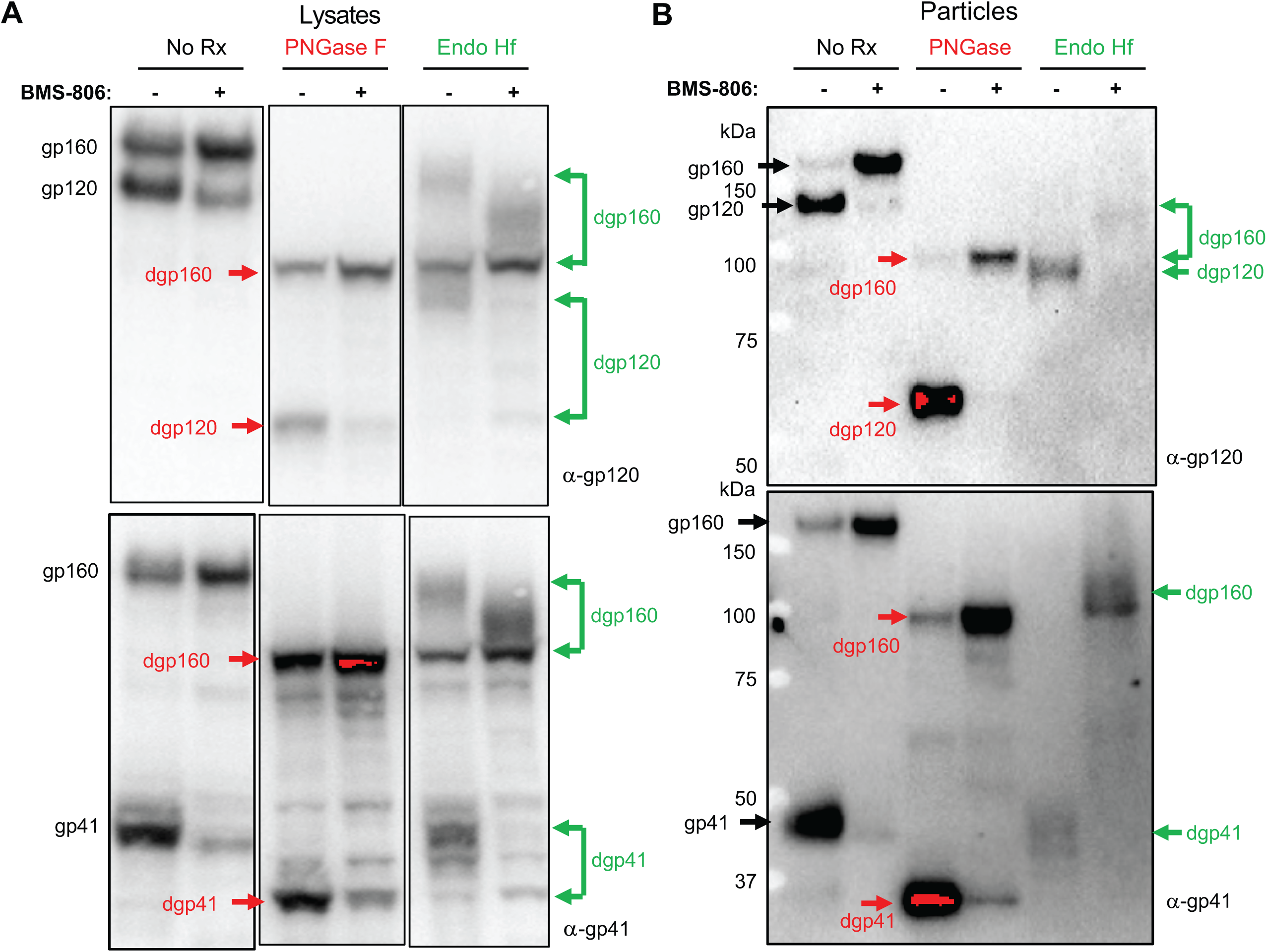
Effect of BMS-806 on the synthesis, processing and glycosylation of wild-type HIV-1_AD8_ Env. A549-Gag/Env cells were treated with BMS-806 (10 µM) or mock treated during doxycycline induction of Gag/Env expression. Lysates were prepared from cells (A) and supernatants containing virus-like particles (VLPs) (B), and were treated with Peptide-N-glycosidase F (PNGase F) or Endoglycosidase Hf (Endo Hf), or mock treated (no Rx). The Envs were run on reducing SDS-polyacrylamide gels and analyzed by Western blotting. The deglycosylated gp160, gp120 and gp41 proteins (dgp160, dgp120 and dgp41, respectively) are indicated by arrows (red – PNGase F-treated sample; green – Endo Hf-treated sample). Data in this figure are representative of those obtained in two independent experiments.

## DISCUSSION

The uncleaved HIV-1 Env serves as a precursor to the cleaved functional Env and, by eliciting poorly neutralizing antibodies, as a potential decoy to the host immune system. Antibody or ligand binding and smFRET analyses indicate that the Env precursor can sample multiple conformations that resemble States 1, 2 and 3 of the mature viral Env spike (73–81). The conformational plasticity of the Env precursor contrasts with the behavior of the mature Env, which in the absence of ligands largely resides in State 1 (42, 81). Therefore, proteolytic cleavage stabilizes State-1 Env, which is highly resistant to neutralization by antibodies recognizing other Env conformations. Although proteolytic maturation also primes the membrane-fusing potential of other Class I viral membrane fusion proteins, the effects of cleavage on HIV-1 Env conformational plasticity are unusual. For example, crystal structures comparing the influenza virus precursor, HA0, with the cleaved HA1/HA2 trimer showed differences only in the immediate vicinity of the cleavage site (120). Uncleaved HIV-1 Envs can be transported from the endoplasmic reticulum to the cell surface by bypassing the Golgi or, when trafficking through the classical secretory pathway, by escaping furin cleavage in the Golgi (72). Both subsets of uncleaved Envs on the surface of expressing cells can be recognized by pNAbs and therefore represent a potentially abundant source of Env conformations other than State 1 (72, 79). The resulting diversion of host antibody responses away from State-1 Env, the major target for neutralizing antibodies, would have considerable advantages for a persistent virus like HIV-1.

BMS-806 can enrich State 1 in the uncleaved membrane-anchored Env (79, 81) and BS3 crosslinking could hypothetically help to stabilize this conformation. Nonetheless, once Env(-) glycoproteins were solubilized in detergent, these treatments did not prevent Env(-) from assuming non-State-1 conformations. The loss of membrane interactions (122), the effects of detergents or other manipulations during purification may have contributed to diminished State-1 occupancy in this case.

**Our structural and biophysical analyses indicate that the cleaved Env conformation seen in the sgp140 SOSIP.664 trimers is also sampled by the uncleaved Env, but notably, in an asymmetric fashion. Thus, a**lthough the asymmetry of the U_1_ and U_2_ uncleaved Env trimers alters the quaternary relationships among the Env protomers, the fold of the individual Env(-) protomers resembles those of sgp140 SOSIP.664 and PGT151-bound EnvΔCT trimers. Analysis by smFRET has suggested that these Envs are predominantly in a State-2-like conformation (121). By analogy, we deduce that U_1_ and U_2_ represent State-2-like conformations. State 2 has been suggested to represent a default intermediate conformation favored by Envs that experience a destabilization of State 1 (48,49,54,82,121). CD4 binding to the wild-type HIV-1 Env trimer sequentially induces State-2 and State-3 conformations in the bound protomer, whereas the other, ligand-free protomers in the Env trimer assume State-2 conformations (49). Although PGT151 is a broadly neutralizing antibody and can presumably interact with State-1 Envs, it induces asymmetry in the Env trimer, causing the Env protomers to assume State-2-like conformations (121). Thus, breaking symmetry in the HIV-1 Env trimer often results in the adoption of a State-2 conformation, consistent with the proposed default nature of this intermediate.

**Asymmetry of both uncleaved and cleaved Env trimers appears to be related to the structural plasticity and flexibility of the gp41 HR1_N_ region, which is directly situated in the inter-protomer interface and is allosterically coupled with the quaternary Env conformation. On the one hand, the HR1_N_ structure can directly affect the packing of the central 3-HB_C_ coiled coils; on the other hand, the HR1_N_ rigidity can allosterically regulate the inter-protomer opening angle. Mutagenesis studies have suggested that in the pre-triggered (State-1) Env conformation, the HR1_N_ region contributes to the non-covalent association of gp120 with gp41 (109-111). We observed a relationship between the inter-protomer opening angle of asymmetric Env trimers and HR1_N_ helicity. As initial CD4 binding to the Env trimer occurs asymmetrically, with State-2 conformations assumed by the unbound protomers** (**49**)**, the HR1_N_ regions presumably transition from as-yet-unknown State-1 conformations to predominantly helical conformations. Subsequent assembly of three HR1_N_ helices into the extended gp41 coiled coil [(HR1_N+C_)_3_] projects the fusion peptide toward the target membrane.**

**The symmetry of the mature, pre-triggered (State-1) HIV-1 Env trimer likely contributes to its ability to evade pNAbs. Supporting this assertion is the previous observation that the fraction of cell-surface Env recognized by bNAbs crosslinked into trimers, whereas the cell-surface Env that was recognized by pNAbs crosslinked into dimers and monomers, possible reflecting trimer asymmetry** (**72**)**. The asymmetry observed for the uncleaved Env**(**-**) **U_1_ and U_2_ trimers potentially allows pNAbs to access their epitopes with minimal steric hindrance. Indeed, pNAbs directed against the gp120 V3 region or CD4-binding site can be docked into the open face of the U_1_ Env trimer with only minimal readjustment of surrounding structures to remove steric clashes (data not shown). Maintaining C3 symmetry may be one prerequisite for preserving an antibody-resistant State-1 Env conformation. Our study implicates the conformationally labile gp41 HR1_N_ segment in maintaining trimer symmetry, and the high-resolution structure of this functionally important region in a State-1-compatible Env conformation is a future goal.**

The intrinsic conformational heterogeneity of the uncleaved HIV-1 Env trimer and the low occupancy of certain conformational states present significant challenges to their structural characterization. Previous studies of detergent-solubilized uncleaved HIV-1 Envs with truncated cytoplasmic tails were performed without extensive 3D classification and with C3 symmetry imposed, resulting in lower-resolution structures (123, 124). **Our current study takes advantage of subsequent advances in 3D classification in cryo-EM technology and data processing to identify two major classes of Env**(-) **trimers, both asymmetric. Cryo-EM and smFRET analyses support the existence of other conformations in the Env**(-) **preparation, but high-resolution reconstruction of these conformers was unsuccessful** (**95**)**. Current 3D hierarchical classification methods are prone to ignore or completely miss lowly populated conformational states or experience difficulties in precisely classifying these low-population conformations, which then leads to insufficient resolution for structure determination and functional interpretation** (**125**)**. A more complete characterization of the multiple conformations assumed by the uncleaved HIV-1 Env may require approaches better able to deal with a high degree of structural heterogeneity than maximum-likelihood-based 3D classification** (**125,126**).

BMS-806 inhibits HIV-1 entry, blocking CD4-induced transitions of the mature Env from a pre-triggered (State-1) conformation to downstream states (42,53,54,79,81). On the cell or viral membrane, uncleaved Env can respond to treatment with BMS-806 by increasing the occupancy of State 1 (79, 81). Consequently, BMS-806 decreases recognition of uncleaved Env by pNAbs, whereas recognition by most bNAbs is maintained or increased (55, 79). We found that BMS-806 also exerts a significant effect on Env during its maturation. BMS-806 treatment of cells expressing wild-type HIV-1 Env resulted in decreases in both gp160 cleavage and modification by complex carbohydrate structures; transport through the Golgi and incorporation into VLPs were not apparently blocked by BMS-806. These observations imply that gp160 conformational flexibility contributes to the efficiency with which the Env precursor is acted upon by furin and glycosylation enzymes. The requirement that functional Env is cleaved (25, 127) therefore provides selective pressure to maintain flexibility in the HIV-1 Env precursor. The resulting conformational heterogeneity of the Env precursor represents a potential advantage for a persistent virus like HIV-1 by skewing host antibody responses away from State 1. For immunization strategies employing membrane-anchored HIV-1 Env or during natural HIV-1 infection, treatment with BMS-806 analogues could potentially increase the presentation of the State-1 Env conformation to the immune system. BMS-806 analogues (79) could also assist future investigation of State-1-like conformations of uncleaved and cleaved HIV-1 Env trimers.

## MATERIALS AND METHODS

### Protein expression and purification

For expression of the uncleaved full-length membrane-anchored HIV-1_JR-FL_ Env(-) glycoprotein, the env cDNA was codon-optimized and was cloned into an HIV-1-based lentiviral vector. These Env sequences contain a heterologous signal sequence from CD5 in place of that of wild-type HIV-1 Env. The proteolytic cleavage site between gp120 and gp41 was altered, substituting two serine residues for Arg 508 and Arg 511. In the HIV-1_JR-FL_ Env(-) glycoprotein, the amino acid sequence LVPRGS-(His)_6_ was added to the C-terminus of the cytoplasmic tail. For Env(-) expression, the *env* coding sequences were cloned immediately downstream of the tetracycline (Tet)-responsive element (TRE). Our expression strategy further incorporated an internal ribosomal entry site (IRES) and a contiguous puromycin (puro) T2A enhanced green fluorescent protein (EGFP) open reading frame downstream of env (TRE-*env*-IRES-puro.T2A.EGFP). Uncleaved membrane-anchored Env(-) was produced by exogenous expression in CHO cells. Briefly, the HIV-1-based lentiviral vector encoding HIV-1_JR-FL_ Env(-) was packaged, pseudotyped with the vesicular stomatitis virus (VSV) G protein, and used to transduce CHO cells (Invitrogen) constitutively expressing the reverse Tet transactivator (rtTA). High-producer clonal cell lines were derived using a FACSAria cell sorter (BD Biosciences) to isolate individual cells expressing high levels of EGFP. The integrity of the recombinant env sequence in the clonal lines was confirmed by sequence analysis of PCR amplicons. Clonal cultures were adapted for growth in a serum-free suspension culture medium (CDM4CHO; Thermo Fisher).

For the exogenous production of the Env(-) glycoprotein, cells were expanded in a suspension culture using a roller bottle system (Thermo) and were treated with 1 μg/ml of doxycycline and 10 μM BMS-378806 (herein referred to as BMS-806) (Selleckchem) after reaching a density of >4 × 10^6^ cells per ml. After 18 to 24 h of culture with doxycycline and BMS-806, the cells were harvested by centrifugation. During the remainder of the purification procedure, 10 μM BMS-806 was added to all buffers. The cell pellets were homogenized in a homogenization buffer (250 mM sucrose, 10 mM Tris-HCl [pH 7.4], and a cocktail of protease inhibitors [Roche Complete EDTA-free tablets]). Membranes were then extracted from the homogenates by ultracentrifugation. The extracted crude membrane pellet was collected, resuspended in 1×PBS to a final concentration of 5 mg of wet membrane per ml of 1×PBS and crosslinked with 5 mM BS3 (Proteochem), followed by solubilization with a solubilization buffer containing 100 mM (NH_4_)_2_SO_4_, 20 mM Tris-HCl (pH 8.0), 300 mM NaCl, 20 mM imidazole, 1% (wt/vol) Cymal-5 (Anatrace), and a cocktail of protease inhibitors (Roche Complete EDTA-free tablets). The suspension was ultracentrifuged for 30 min at 100,000 × g and 4°C. The supernatant was collected and was mixed with a small volume of preequilibrated Ni-nitrilotriacetic acid (NTA) beads (Qiagen) for 2 h on a rocking platform at 4°C. The mixture was then injected into a small column and washed with a buffer containing 20 mM Tris-HCl (pH 8.0), 100 mM (NH_4_)_2_SO_4_, 1 M NaCl, 30 mM imidazole, and 0.5% Cymal-5. The beads were resuspended in a buffer containing 20 mM Tris-HCl (pH 8.0), 100 mM (NH_4_)_2_SO_4_, 250 mM NaCl, 4.5 mg/ml Amphipol A8-35 (Anatrace), 0.006% DMNG (Anatrace) and a cocktail of protease inhibitors (Roche Complete EDTA-free tablets), and incubated for 2 hours on a rocking platform. The mixture was applied to a column and the buffer was allowed to flow through. The beads were then resuspended in a buffer containing 20 mM Tris-HCl (pH 8.0), 100 mM (NH_4_)_2_SO_4_, 250 mM NaCl, 4.5 mg/ml Amphipol A8-35 (Anatrace) and a cocktail of protease inhibitors (Roche Complete EDTA-free tablets), and incubated for 2 hours on a rocking platform. The mixture was added to a column and the buffer allowed to flow through, followed by washing with 10 bed volumes of a buffer containing 20 mM Tris-HCl (pH 8.0), 100 mM (NH_4_)_2_SO_4_, and 250 mM NaCl. Proteins were eluted from the bead-filled column with a buffer containing 20 mM Tris-HCl (pH 8.0), 100 mM (NH_4_)_2_SO_4_, 250 mM NaCl, and 250 mM imidazole. The buffer of the eluted Env(-) glycoprotein solution was exchanged with imaging buffer containing 20 mM Tris-HCl (pH 8.0), 100 mM (NH_4_)_2_SO_4_, and 250 mM NaCl with a Centrifugal Filter (Millipore), and was concentrated. Before cryo-plunging, Cymal-6 (Anatrace) was added to the Env(-) glycoprotein solution at a final concentration of 0.005%.

### Expression of wild-type HIV-1 Env and virus-like particles (VLPs)

Human A549 lung epithelial cells (ATCC) inducibly expressing Env and an HIV-1 Gag-mCherry fusion protein under the control of a tetracycline-regulated promoter were established as described (72). Briefly, A549-rtTA cells constitutively expressing the reverse tet transactivator were transduced with an HIV-1-based lentivirus vector expressing Rev and Env from HIV-1_AD8_, a primary HIV-1 strain (128). These A549-Env cells were transduced with a lentivirus vector expressing the HIV-1 Gag precursor fused with mCherry (72). The doxycycline-regulated expression of the Gag-mCherry fusion protein resulted in the release of Env-containing VLPs into the medium. Herein, we designate these cells A549-Gag/Env. The A549-Gag/Env cells were grown in DMEM/F12 supplemented with 10% FBS, L-glutamine and penicillin-streptomycin.

### Antibodies

Antibodies against HIV-1 Env were kindly supplied by Dr. Dennis Burton (Scripps), Drs. Peter Kwong and John Mascola (Vaccine Research Center, NIH), Dr. Barton Haynes (Duke), Dr. Hermann Katinger (Polymun), Dr. James Robinson (Tulane) and Dr. Marshall Posner (Mount Sinai Medical Center). In some cases, anti-Env antibodies were obtained through the NIH AIDS Reagent Program. Antibodies for Western blotting include goat anti-gp120 polyclonal antibody (ThermoFisher) and the 4E10 human anti-gp41 antibody (Polymun). An HRP-conjugated goat anti-human IgG (Santa Cruz) and an HRP-conjugated goat anti-rabbit antibody (Santa Cruz) were used as secondary antibodies for Western blotting.

### Single-molecule FRET: sample preparation, data acquisition and analysis

Analysis of the conformational dynamics of HIV-1 Env was performed after enzymatic labeling of the V1 and V4 regions of gp120 on the purified (His)_6_-tagged HIV-1_JR-FL_ Env(-) glycoprotein with Cy3 and Cy5 fluorophores, respectively, as previously described (42). A transfection ratio of 20:1 of non-tagged: V1/V4-tagged HIV-1_JR-FL_ Env(-) was used to ensure that only one protomer within a trimer carries enzymatic tags for site-specific labeling. The HIV-1_JR-FL_ Env(-) glycoprotein was purified from transiently expressing 293T cells that had been treated with BMS-806 and crosslinked with BS3, as described above. The purified HIV-1_JR-FL_ Env(-) glycoprotein in buffer (20 mM Tris-HCl (pH 8.0), 10 mM MgCl_2_, 10 mM CaCl_2_, 100 mM (NH4)_2_SO_4_, 250 mM NaCl, 0.005% Cymal-6, 10 µM BMS-806) was labeled with Cy3B(3S)-cadaverine (0.5 µM), transglutaminase (0.65 µM; Sigma Aldrich), LD650-CoA (0.5 µM) (Lumidyne Technologies), and AcpS (5 µM) at room temperature overnight. After labeling, Env(-) trimers were purified using Zeba^TM^ spin desalting columns (ThermoFisher) to remove free dyes. Finally, prior to imaging, fluorescence-labeled HIV-1_JR-FL_ Env(-) carrying the (His)_6_ epitope tag was incubated with biotin-conjugated anti-(His)_6_ tag antibody (HIS.H8, Invitrogen) at 4° for two hours.

All smFRET data were acquired on a home-built total internal reflection fluorescence (TIRF) microscope, as previously described (42, 129). Fluorescently labeled HIV-1_JR-FL_ Env(-) trimers were immobilized on passivated streptavidin-coated quartz microscopy slides and washed with pre-imaging buffer specifically made for this experiment. The pre-imaging buffer consisted of 20 mM Tris HCl (pH 8.0), 100 mM (NH4)_2_SO_4_, 250 mM NaCl, 0.005% Cymal-6, and 10 µM BMS-806. For smFRET analysis, a cocktail of triplet-state quenchers and 2 mM protocatechuic acid (PCA) with 8 nM protocatechuic 3,4-dioxygenase (PCD) were added to the above pre-imaging buffer to remove molecular oxygen. Cy3 and Cy5 fluorescence was detected with a 60x water-immersion objective (Nikon), split by a diachronic mirror (Chroma), and imaged on two synchronized ORCA-Flash4.0v2 sCMOS cameras (Hamamatsu) at 40 frames/seconds for 80 seconds.

smFRET data analysis was performed on the customized Matlab (Mathworks) program SPARTAN (129). Fluorescence intensity trajectories were extracted from recorded movies, and FRET efficiency (FRET) was calculated based on FRET= I_A_/(ϒI_D_+I_A_), where I_D_ and I_A_ are the fluorescence intensities of donor (D) and acceptor (A), respectively, and ϒ is the correlation coefficient, which incorporates the difference in quantum yields of donor and acceptor and detection efficiencies of the donor and acceptor channels. FRET trajectories were further compiled into a FRET histogram, which provides information about the distribution of Env(-) molecules among the conformational states. The state distributions in the FRET histogram were then fitted to the sum of three Gaussian distributions (based on previously identified FRET trajectories) (42,81,121) in Matlab, and the occupancy of each state was further obtained from the area under each Gaussian distribution.

### Immunoprecipitation of cell-surface Env

One day prior to transfection, HOS cells were seeded in 6-well plates (6 x 10^5^ cells/well). The cells were transfected the next day with 0.4 µg of the pSVIIIenv plasmid expressing the wild-type HIV-1_JR-FL_ Env and 0.05 µg of a Tat-expressing plasmid. Two days later, the cells were washed twice with blocking buffer (1×PBS with 5% FBS) and then incubated for 1 hour at 4°C with 6 µg/µl anti-gp120 monoclonal antibody. Cells were then washed four times with blocking buffer, four times with washing buffer (140 mM NaCl, 1.8 mM CaCl_2_, 1 mM MgCl_2_ and 20 mM Tris, pH 7.5), and lysed in NP-40 buffer (0.5 % NP-40, 0.5 M NaCl and 10 mM Tris, pH 7.5) for 5 min at 4°C with gentle agitation. Lysates were cleared by centrifugation at 15,000 x g for 30 min at 4°C. Antibody-bound Env was precipitated using Protein A-Sepharose beads and analyzed by SDS-PAGE and Western blotting with a horseradish peroxidase (HRP)-conjugated rabbit anti-gp120 polyclonal serum.

### Cell-based enzyme-linked immunosorbent assay (ELISA)

CHO cells expressing HIV-1_JR-FL_ Env(-) were induced with 1 µg/ml doxycycline with or without 10 µM BMS-806. Fifteen to twenty-four hours later, the cells were washed twice with washing buffer #1 (20 mM Hepes, pH 7.5, 1.8 mM CaCl_2_, 1 mM MgCl_2_, 140 mM NaCl), and crosslinked with 5 mM BS3 or incubated in buffer without crosslinker. Forty-five minutes later, the cells were quenched with quench buffer (50 mM Tris, pH 8.0, 1.8 mM CaCl_2_, 1 mM MgCl_2_, 140 mM NaCl). The cells were blocked with a blocking buffer (35 mg/ml BSA, 10 mg/ml non-fat dry milk, 1.8 mM CaCl_2_, 1 mM MgCl_2_, 25 mM Tris, pH 7.5 and 140 mM NaCl) and incubated with the indicated primary antibody in blocking buffer for 30 min at 37°C. Cells were then washed three times with blocking buffer and three times with washing buffer #2 (140 mM NaCl, 1.8 mM CaCl_2_, 1 mM MgCl_2_ and 20 mM Tris, pH 7.5) and re-blocked with the blocking buffer. A horseradish peroxidase (HRP)-conjugated antibody specific for the Fc region of human IgG was then incubated with the samples for 45 min at room temperature. Cells were washed three times with blocking buffer and three times with washing buffer #2. HRP enzyme activity was determined after addition of 35 µl per well of a 1:1 mix of Western Lightning oxidizing and luminal reagents (Perkin Elmer Life Sciences) supplemented with 150 mM NaCl. Light emission was measured with a Mithras LB940 luminometer (Berthold Technologies).

### Analysis of Env(-) glycoforms in BMS-806-treated cells

CHO cells expressing HIV-1_JR-FL_ Env(-) were treated with 1 µM BMS-806 or an equivalent volume of the carrier, DMSO. After 18-24 h of culture, the cells were harvested and lysed in homogenization buffer (see above) and treated with different glycosidases following the manufacturer’s instructions. The lysates were analyzed by Western blotting with a horseradish peroxidase (HRP)-conjugated anti-HIV-1 gp120 antibody, as described above.

### Analysis of Env glycopeptides

The sample preparation and mass spectrometric analysis of Env(-) glycopeptides has been described previously (39, 40), and no changes were made to the procedure for the current analysis. Briefly, the Env(-) glycoprotein was denatured with urea, reduced with TCEP, alkylated with iodoacetamide, and quenched with dithiothreitol. The protein was then buffer exchanged, digested with trypsin alone or with a combination of trypsin and chymotrypsin, generating glycopeptides.

The glycopeptides were analyzed by LC-MS on an LTQ-Orbitrap Velos Pro (Thermo Scientific) mass spectrometer equipped with ETD (electron transfer dissociation) that was coupled to an Acquity Ultra Performance Liquid Chromatography (UPLC) system (Waters). About 35 micromoles of digest was separated by reverse phase HPLC using a multistep gradient, on a C18 PepMap™ 300 column. The mass spectrometric analysis was performed using data-dependent scanning, alternating a high-resolution scan (30,000 at m/z 400), followed by ETD and collision-induced dissociation (CID) data of the five most intense ions. The glycopeptides were identified in the raw data files using a combination of freely available glycopeptide analysis software and expert identification, as described previously (39).

### Analysis of A549-Gag/Env cells and VLPs treated with BMS-806

To analyze the effect of BMS-806 on the processing of the wild-type HIV-1_AD8_ Env, 150-mm dishes of 30-40% confluent A549-Gag/Env cells were seeded and, on the following day, treated with 2 µg/ml doxycycline. At the same time, 10 µM BMS-806 was added. Approximately 72 hours after induction, cell lysates and medium were harvested. To prepare VLPs, the culture medium was cleared by low-speed centrifugation (500 x g for 15 minutes at 4°C) and 0.45-µm filtration. VLPs were pelleted by centrifugation at 100,000 x g for one hour at 4°C. The resuspended VLP preparation was clarified by low-speed centrifugation.

Env solubilized from cell lysates and VLPs was denatured by boiling in denaturing buffer (New England Biolabs) for 10 minutes. Samples were mock-treated or treated with PNGaseF or Endo Hf (New England Biolabs) for 1.5 hours according to the manufacturer’s protocol. The treated samples were then analyzed by reducing SDS-PAGE and Western blotting.

### Cryo-EM sample preparation

A 3-μl drop of 0.3 mg/ml Env(-) protein solution was applied to a glow-discharged C-flat grid (R1/1 and R1.2/1.3, 400 Mesh, Protochips, CA, USA), blotted for 2 sec, then plunged into liquid ethane and flash-frozen using an FEI Vitrobot Mark IV.

### Cryo-EM data collection

Cryo-EM grids were first visually screened on a Tecnai Arctica transmission electron microscope (FEI) operating at 200 kV. Qualified grids were then imaged in a 200-kV FEI Tecnai Arctica microscope, equipped with an Autoloader, at a nominal magnification of 210,000 times, and in a 300-kV Titan Krios electron microscope (FEI) equipped with a Gatan BioQuantum energy filter, at a nominal magnification of 105,000 times, operating at 300 kV. Coma-free alignment and astigmatism were manually optimized prior to data collection. Cryo-EM data from the 200-kV Arctica microscope were collected semi-automatically by Leginon version 3.1 (130, 131) on the Gatan K2 Summit direct electron detector camera (Gatan Inc., CA, USA) in a super-resolution counting mode, with a dose rate of 8 electrons/pixel/second and an accumulated dose of 50 electrons/Å^2^ over 38 frames per movie. The calibrated physical pixel size and the super-resolution pixel size were 1.52 Å and 0.76 Å, respectively. The defocus for data collection was set in the range of -1.0 to -3.0 µm. A total of 12,440 movies were collected on the 200-kV Arctica microscope without tilting the stage, from which 10,299 movies were selected for further data analysis after screening and inspection of data quality.

Cryo-EM data from the 300-kV Krios microscope, including both zero-tilted and 45°-tilted images, were collected on the K2 Summit direct electron detector (Gatan) at a pixel size of 0.685 Å in a super-resolution counting mode, with an accumulated dose of ∼53 electrons/Å^2^ across 40 frames per movie. With defocus ranging from -1.0 to -2.7 μm, a total of 10,929 movies were acquired across three sessions.

Zero-tilted and 45°-tilted images were collected by a semi-automatic process set up in Serial EM (132), which is compatible with customized scripts. For the collection of zero-tilted movies, the process normally involved the following steps: Square selection and focusing, hole selection, serial local focusing and data acquisition. In the final step, precise adjustment of the defocus was conducted each time before recording movies for a new group of holes. However, for the collection of tilted movies, precise adjustment of the defocus was performed for all holes in the first place, followed by an extra coordinate transformation for the x-axis and y-axis. Tilted movies were then recorded serially with the new defocus and coordinates.

### Cryo-EM data processing and analysis

The raw movie frames of each dataset were first corrected for their gain reference and each movie was used to generate a micrograph that was corrected for sample movement and drift with the MotionCor2 program (133) at a super-resolution pixel size (0.76 Å for the 200-kV dataset, 0.685 Å for the 300-kV dataset). These drift-corrected micrographs were used for the determination of the actual defocus of each micrograph with the CTFFind4 (134) and Gctf (135) programs. Icy or damaged micrographs were removed through manual per-image screening.

For the 200-kV dataset, using DeepEM, a deep learning-based particle extraction program that we developed (136), 1,436,424 particles of Env(-) were automatically selected in a template-free fashion. All 2D and 3D classifications were done at a pixel size of 1.52 Å. After the first round of reference-free 2D classification, bad particles were rejected upon inspection of class-average quality, which left 1,366,095 particles. The initial model, low-pass filtered to 60 Å, was used as the input reference to conduct unsupervised 3D classification into 5 classes with C3 symmetry, using an angular sampling of 7.5° and a regularization parameter T of 4. Iterative 3D classification in RELION (137) and ROME (138) resulted in a 3D class of 121,979 particles that reached a resolution of 5.5 Å (gold-standard FSC at 0.143 cutoff) after refinement, with imposition of C3 symmetry. More details of this preliminary, intermediate analysis were described in an online bioRxiv preprint (95).

For the zero-tilt 300-kV dataset, micrographs without dose-weighting were used by Gctf (135) to estimate the global CTF parameters; for the 45°-tilt dataset, particles were first picked by a program based on a VGG deep neural network improved from the DeepEM algorithm design (136). The coordinates were then applied for local CTF estimation in Gctf (135). We found that for most of 45°-tilted micrographs, limiting the resolution range used for CTF determination in Fourier space improved the accuracy of the resulting CTF parameters. This was realized by including the variables “local_resL” and “local_resH” in the Gctf (135) command. Automatic picking followed by manual examination yielded 1,941,541 particles of the HIV-1_JR-FL_ Env(-) trimers, with 785,844 zero-tilted and 1,155,697 tilted particles.

All 2D and 3D classifications of the particles from the 300-kV datasets were conducted with dose-weighted micrographs generated by MotionCor2 (133). Particles were stacked at 2.74 Å/pixel using a box size of 84*84 for initial sorting. Two rounds of reference-free 2D classification were performed in RELION 3.0 (137), followed by one round in ROME (138), which combines maximum likelihood-based image alignment and statistical manifold learning-based classification. Bad particles were rejected upon inspection of the class average’s quality after each round of 2D classification, leaving 572,205 particles for 3D refinement. The initial model was generated in RELION 3.0 (137) using particles from diversely oriented 2D classes, and was low-pass filtered to 60 Å.

3D classification and refinement of the 300-kV datatset were performed in RELION 3.0 (137), as summarized in Table 3. In the first round of unsupervised 3D classification, the Healpix order was enhanced from 2 to 3 at the 20th iteration. To prevent tilted particles from being separated as a sole 3D class, the resolution limit to restrict the probability calculation was set at 15 Å in the preceding 20 iterations and 10 Å in the posterior iterations. The 2nd round of 3D classification retained the same parameters except that K (the number of classes) was changed to 6. The 3rd round of 3D classification was performed by local searching (σ=4, meaning that the standard deviation of the Euler angles equals 4 times the Healpix order) to discard amorphous particles. Particles with the correct size and detailed secondary structures were selected and binned two-fold into 1.37 Å/pixel for further refinement. The selected 278,582 particles were first aligned together by auto-refinement, and then were classified into 12 classes within a soft, global mask without alignment. Particles from 5 classes with complete domain constitution were sorted out and used for per-particle CTF refinement in RELION 3.0 (137). Imposed with updated CTF correction, the sorted stacks were classified with local searching into two major classes.

**Table 2.** Cryo-EM data collection, refinement and validation statistics.

|  | HIV-1<sub>JR-FL</sub><br>Env(-) U<sub>1</sub> | HIV-1<sub>JR-FL</sub><br>Env(-) U<sub>2</sub> |
| --- | --- | --- |
| **Data collection and processing** |  |  |
| Magnification | 105,000 | 105,000 |
| Voltage (kV) | 300 | 300 |
| Electron exposure (e/Å) | 53 | 53 |
| Defocus range (μm) | -1.0 to -2.7 | -1.0 to -2.7 |
| Pixel size (Å) | 0.685 | 0.685 |
| Symmetry imposed | C1 | C1 |
| Initial particle images (no.) | 572,205 | 572,205 |
| Final particle images (no.) | 123,372 | 55,571 |
| Map resolution (Å) | 4.1 | 4.7 |
| FSC threshold | 0.143 | 0.143 |
| Map resolution range (Å) | 3.8 to 8 | 3.8 to 10 |
| **Refinement** |  |  |
| Initial model used (PDB code) | 5FUU | 5FUU |
| Model resolution (Å) | 4.2 | 4.8 |
| FSC threshold | 0.143 | 0.143 |
| Model resolution range (Å) | 3.8 to 8 | 3.8 to 10 |
| Map sharpening B factor (Å<sup>2</sup>) | -75 | -75 |
| Model composition |  |  |
| Non-hydrogen atoms | 14070 | 14062 |
| Protein residues | 1776 | 1775 |
| Ligands | 3 | 3 |
| B factors (Å<sup>2</sup>) |  |  |
| Protein | 191.26 | 191.13 |
| Ligands | 3.16 | 12.91 |
| **R.m.s. deviations** |  |  |
| Bond lengths (Å) | 0.008 | 0.008 |
| Bond angles (degree) | 1.455 | 1.204 |
| **Validation** |  |  |
| MolProbity score | 2.67 | 2.86 |
| Clashscore | 34.63 | 46.01 |
| Poor rotamers (%) | 0.18 | 1.78 |
| **Ramachandran plot** |  |  |
| Favored (%) | 85.97 | 90.65 |
| Allowed (%) | 13.75 | 9.06 |
| Disallowed (%) | 0.29 | 0.29 |

**Table 3.** Summary of 3D classification parameters (300-kV dataset)

| Iteration number | K | Healpix order | Global searching<br>or local<br>searching | Particles left for<br>next round |
| --- | --- | --- | --- | --- |
| 1 | 4 | 2 & 3 | Global | 479,120 |
| 2 | 6 | 2 & 3 | Global | 362,017 |
| 3 | 8 | 4 | Local, $\sigma=4$ | 278,582 |
| 4 | 12 | — | — | 271,277 |
| 5 | 8 | 4 | Local, $\sigma=4$ | 243,313 |
| *Retrieve and Combine* |  |  |  |  |
| 6 | 4 | 2 & 3 | Global | 284,664 |
| 7 | 3 | 2 | Global | 269,801 |
| 8 | 4 | 2 | Global | 265,901 |
| 9 | 8 | 4 | Local, $\sigma=4$ | 229,246 |
| 10 | 6 | 4 | Local, $\sigma=4$ | 223,613 |
| 11 | 6 | 4 | Local, $\sigma=8$ | 211,023 |
| 12 | 6 | 4 | Local, $\sigma=4$ | 164,789 |
| 13 | 5 | 5 | Local, $\sigma=4$ | 156,714 |
| 14 | 5 | 5 | Local, $\sigma=4$ | — |

As observed in Chimera (139), the distribution of particles concentrated in the top-view orientation for both maps, leading to anisotropy of the final resolution. Therefore, we retrieved the tilt-view particles excluded by previous rounds of 3D classification, and combined them with particles from the two classes. This was accomplished by several rounds of screening satisfying classes from the results of deep 2D classification in ROME (138). The new particle dataset, containing 171,342 zero-tilted particles and 157,607 45°-tilted particles, was used for one round of 3D classification under global searching with Healpix order 2. Particles from 3 of the 4 classes were identified as HIV-1_JR-FL_ Env(-) trimers with improved isotropic resolution; these 284,664 particles were combined for the next round of 3D classification. Another round of 3D classification using the same parameters except for K=3 was performed to exclude particles with poor quality. The principal class consisting of 92% of this round’s particles was reserved.

For elaborate 3D classification, we adopted a hierarchical enhancement of Healpix order in the next 9 rounds (Table 3): Sorted particles from the previous round of 3D classification were used for auto-refinement followed by classification into four classes with local searching under a Healpix order of 4. In every round, this process produced a major class consistent with the structure of the conventional Env trimer and consisting of more than 80% of the input particles, while the other classes appeared in incomplete form. Therefore, this major class of particles was used for auto-refinement and was chosen as input for next round of 3D classification. This classification-selection-refinement-classification process was iterated four times, using different K (class number) values and the same Healpix order 4, until the result demonstrated more than one principal class. C1 symmetry was imposed throughout all these unsupervised 3D classifications. In the last two rounds, we enhanced the Healpix order to 5 to perform local-searching 3D classification again, and finally obtained five classes. Four of these classes, consisting of 96% of the input particles, exhibited different degrees of asymmetry. By carefully comparing their features, two classes with similar topology were designated State-U_1_ while the other two classes were designated State-U_2_, containing 123,372 and 55,571 particles respectively. The last round of auto-refinement for the U_1_ and U_2_ datasets was done in RELION 3.0 (137), applied with a soft-edged global mask when it fell into local searching. According to the in-plane shift and Euler angles of each particle from the final refinement, we reconstructed the two half-maps of each state at a super-resolution counting mode with a pixel size of 0.685 Å. The overall masked resolutions for the reconstructed maps of State-U_1_ and State-U_2_ were 4.1 Å and 4.7 Å respectively, measured by the gold-standard FSC at 0.143-cutoff.

### Atomic model building and refinement

The symmetric structure of the HIV-1_BG505_ sgp140 SOSIP.664 trimer with three BMS-806 molecules bound (PDB: 6MTJ) (118) and the asymmetric structure of the HIV-1_JR-FL_ EnvΔCT glycoprotein bound to PGT151 Fabs (PDB: 5FUU) (105) were used as reference models to build a U_1_ structure. The template structures were docked in Coot (140), and then main-chain and side-chain fitting was improved manually to generate the starting coordinate file. The fitting of the U_1_ model was further improved by real_space_refinement with secondary structure restraints in Phenix (141). Glycans of U_1_ were manually refined in Coot (140) with “Glycan” model, using 5FUU as a reference. The U_1_ model was used as a whole to perform rigid-body fitting into the U_2_ density. Structural comparison was conducted in Pymol (142) and Chimera (139). All figures of the structures were produced in Pymol (142).

### Accession numbers

The cryo-EM reconstructions of states U_1_ and U_2_ reported in this paper have been deposited in the Electron Microscopy Data Bank under accession numbers EMD-XXXX and EMD-XXXX, respectively. The models of U_1_ and U_2_ have been deposited in the Protein Data Bank under ID codes XXXX and XXXX. The cryo-EM raw data, including the motion-corrected micrographs and the particle stacks of U_1_ and U_2_ used for final refinement, have been deposited into the Electron Microscopy Pilot Image Archive (www.ebi.ad.uk/emdb/ampiar) under accession no. EMPIAR-10163.

## Author contributions

J.S. and Y.M. conceived this study. H.Ding and J.C.K. prepared the Env(-)-expressing CHO cells and the A549-Gag/Env cells. S.Z. and R.T.S. analyzed Env(-) antigenicity and established a purification scheme for the Env(-) protein. S.Z. and W.L.W. screened the samples for optimization of cryo-EM imaging. W.L.W. conducted cryo-electron microscopy, collected all data and preprocessed the data. K.W. and S.C. performed data analysis and refined the maps. K.W., S.Z., S.C. and Y.M. built the structural models. E.P.G., S.Z. and H.Desaire analyzed the Env(-) glycans. M.L. and S.Z. conducted smFRET experiments. H.T.N. studied the effect of BMS-806 on the processing of wild-type Env. Y.M. and J.S. wrote the manuscript. All authors contributed to data analysis and manuscript preparation.

## Acknowledgments

This work was funded in part by NIH grants AI125093 (H. Desaire), AI93256, AI100645, AI125093, AI145547, AI127767, AI150471/GM56550 and AI124982 (J.S.), by an Intel academic grant (Y.M.), by grants from the Natural Science Foundation of Beijing Municipality grant No. Z180016/Z18J008 and the National Natural Science Foundation of China grant No. 11774012 (Y.M.), and by a gift from the late William F. McCarty-Cooper. M.L. was supported by a grant (109998-67-RKVA) from the American Foundation for AIDS Research (amfAR). The research was also supported by the Basic Research Core of the University of Alabama, Birmingham Center for AIDS Research (NIH grant AI027767). The cryo-EM experiments were performed in part at the Center for Nanoscale Systems at Harvard University, a member of the National Nanotechnology Coordinated Infrastructure Network (NNCI), which is supported by the National Science Foundation under NSF award no. 1541959. The cryo-EM facility was funded through the NIH grant AI100645, Center for HIV/AIDS Vaccine Immunology and Immunogen Design (CHAVI-ID). The data processing was performed in part in the Sullivan cluster, which is supported by a gift from Mr. and Mrs. Daniel J. Sullivan, Jr.

## REFERENCES

1. Wyatt R, Sodroski J. 1998. The HIV-1 envelope glycoproteins: fusogens, antigens, and immunogens. Science 280:1884–1888.

2. Allan JS, Coligan JE, Barin F, McLane MF, Sodroski JG, Rosen CA, Haseltine WA, Lee TH, Essex M. 1985. Major glycoprotein antigens that induce antibodies in AIDS patients are encoded by HTLV-III. Science 228:1091–1094.

3. Robey WG, Safai B, Oroszlan S, Arthur LO, Gonda MA, Gallo RC, Fischinger PJ. 1985. Characterization of envelope and core structural gene products of HTLV-III with sera from AIDS patients. Science 228:593–595.

4. Klatzmann D, Champagne E, Chamaret S, Gruest J, Guetard D, Hercend T, Gluckman JC, Montagnier L. 1984. T-lymphocyte T4 molecule behaves as the receptor for human retrovirus LAV. Nature 312:767–768.

5. Dalgleish AG, Beverley PC, Clapham PR, Crawford DH, Greaves MF, Weiss RA. 1984. The CD4 (T4) antigen is an essential component of the receptor for the AIDS retrovirus. Nature 312:763–767.

6. Wu L, Gerard NP, Wyatt R, Choe H, Parolin C, Ruffing N, Borsetti A, Cardoso AA, Desjardin E, Newman W, Gerard C, Sodroski J. 1996. CD4-induced interaction of primary HIV-1 gp120 glycoproteins with the chemokine receptor CCR-5. Nature 384:179–183.

7. Trkola A, Dragic T, Arthos J, Binley JM, Olson WC, Allaway GP, Cheng-Mayer C, Robinson J, Maddon PJ, Moore JP. 1996. CD4-dependent, antibody-sensitive interactions between HIV-1 and its co-receptor CCR-5. Nature 384:184–187.

8. Choe H, Farzan M, Sun Y, Sullivan N, Rollins B, Ponath PD, Wu L, Mackay CR, LaRosa G, Newman W, Gerard N, Gerard C, Sodroski J. 1996. The beta-chemokine receptors CCR3 and CCR5 facilitate infection by primary HIV-1 isolates. Cell 85:1135–1148.

9. Deng H, Liu R, Ellmeier W, Choe S, Unutmaz D, Burkhart M, Di Marzio P, Marmon S, Sutton RE, Hill CM, Davis CB, Peiper SC, Schall TJ, Littman DR, Landau NR. 1996. Identification of a major co-receptor for primary isolates of HIV-1. Nature 381:661–666.

10. Dragic T, Litwin V, Allaway GP, Martin SR, Huang Y, Nagashima KA, Cayanan C, Maddon PJ, Koup RA, Moore JP, Paxton WA. 1996. HIV-1 entry into CD4+ cells is mediated by the chemokine receptor CC-CKR-5. Nature 381:667–673.

11. Doranz BJ, Rucker J, Yi Y, Smyth RJ, Samson M, Peiper SC, Parmentier M, Collman RG, Doms RW. 1996. A dual-tropic primary HIV-1 isolate that uses fusin and the beta-chemokine receptors CKR-5, CKR-3, and CKR-2b as fusion cofactors. Cell 85:1149–1158.

12. Feng Y, Broder CC, Kennedy PE, Berger EA. 1996. HIV-1 entry cofactor: functional cDNA cloning of a seven-transmembrane, G protein-coupled receptor. Science 272:872–877.

13. Alkhatib G, Combadiere C, Broder CC, Feng Y, Kennedy PE, Murphy PM, Berger EA. 1996. CC CKR5: a RANTES, MIP-1alpha, MIP-1beta receptor as a fusion cofactor for macrophage-tropic HIV-1. Science 272:1955–1958.

14. Chan DC, Fass D, Berger JM, Kim PS. 1997. Core structure of gp41 from the HIV envelope glycoprotein. Cell 89:263–273.

15. Weissenhorn W, Dessen A, Harrison SC, Skehel JJ, Wiley DC. 1997. Atomic structure of the ectodomain from HIV-1 gp41. Nature 387:426–430.

16. Lu M, Blacklow SC, Kim PS. 1995. A trimeric structural domain of the HIV-1 transmembrane glycoprotein. Nat Struct Biol 2:1075–1082.

17. Karlsson Hedestam GB, Fouchier RA, Phogat S, Burton DR, Sodroski J, Wyatt RT. 2008. The challenges of eliciting neutralizing antibodies to HIV-1 and to influenza virus. Nat Rev Microbiol 6:143–155.

18. Hoxie JA. 2010. Toward an antibody-based HIV-1 vaccine. Annu Rev Med 61:135–152.

19. Haynes BF, Shaw GM, Korber B, Kelsoe G, Sodroski J, Hahn BH, Borrow P, McMichael AJ. 2016. HIV-Host interactions: implications for vaccine design. Cell Host Microbe 19:292–303.

20. Fauci AS. 2016. An HIV vaccine: mapping uncharted territory. JAMA 316:143–144.

21. Fennie C, Lasky LA. 1989. Model for intracellular folding of the human immunodeficiency virus type 1 gp120. J Virol 63:639–646.

22. Li Y, Luo L, Thomas DY, Kang CY. 2000. The HIV-1 Env protein signal sequence retards its cleavage and down-regulates the glycoprotein folding. Virology 272:417–428.

23. Willey RL, Bonifacino JS, Potts BJ, Martin MA, Klausner RD. 1988. Biosynthesis, cleavage, and degradation of the human immunodeficiency virus 1 envelope glycoprotein gp160. Proc Natl Acad Sci U S A 85:9580–9584.

24. Earl PL, Moss B, Doms RW. 1991. Folding, interaction with GRP78-BiP, assembly, and transport of the human immunodeficiency virus type 1 envelope protein. J Virol 65:2047–2055.

25. Bosch V, Pawlita M. 1990. Mutational analysis of the human immunodeficiency virus type 1 env gene product proteolytic cleavage site. J Virol 64:2337–2344.

26. Decroly E, Vandenbranden M, Ruysschaert JM, Cogniaux J, Jacob GS, Howard SC, Marshall G, Kompelli A, Basak A, Jean F, Lazuref C, Bedannet S, Chrétien M, Day R, Seidah NG. 1994. The convertases furin and PC1 can both cleave the human immunodeficiency virus (HIV)-1 envelope glycoprotein gp160 into gp120 (HIV-1 SU) and gp41 (HIV-I TM). J Biol Chem 269:12240–12247.

27. Fenouillet E, Gluckman JC. 1992. Immunological analysis of human immunodeficiency virus type 1 envelope glycoprotein proteolytic cleavage. Virology 187:825–828.

28. Hallenberger S, Bosch V, Angliker H, Shaw E, Klenk HD, Garten W. 1992. Inhibition of furin-mediated cleavage activation of HIV-1 glycoprotein gp160. Nature 360:358–361.

29. Dewar RL, Natarajan V, Vasudevachari MB, Salzman NP. 1989. Synthesis and processing of human immunodeficiency virus type 1 envelope proteins encoded by a recombinant human adenovirus. J Virol 63:129–136.

30. Dewar RL, Vasudevachari MB, Natarajan V, Salzman NP. 1989. Biosynthesis and processing of human immunodeficiency virus type 1 envelope glycoproteins: effects of monensin on glycosylation and transport. J Virol 63:2452–2456.

31. Merkle RK, Helland DE, Welles JL, Shilatifard A, Haseltine WA, Cummings RD. 1991. gp160 of HIV-I synthesized by persistently infected Molt-3 cells is terminally glycosylated: evidence that cleavage of gp160 occurs subsequent to oligosaccharide processing. Arch Biochem Biophys 290:248–257.

32. Kantanen ML, Leinikki P, Kuismanen E. 1995. Endoproteolytic cleavage of HIV-1 gp160 envelope precursor occurs after exit from the trans-Golgi network (TGN). Arch Virol 140:1441–1449.

33. Pfeiffer T, Zentgraf H, Freyaldenhoven B, Bosch V. 1997. Transfer of endoplasmic reticulum and Golgi retention signals to human immunodeficiency virus type 1 gp160 inhibits intracellular transport and proteolytic processing of viral glycoprotein but does not influence the cellular site of virus particle budding. J Gen Virol 78:1745–1753.

34. Miranda L, Wolf J, Pichuantes S, Duke R, Franzusoff A. 1996. Isolation of the human PC6 gene encoding the putative host protease for HIV-1 gp160 processing in CD4+ T lymphocytes. Proc Natl Acad Sci U S A 93:7695–7700.

35. Ohnishi Y, Shioda T, Nakayama K, Iwata S, Gotoh B, Hamaguchi M, Nagai Y. 1994. A furin-defective cell line is able to process correctly the gp160 of human immunodeficiency virus type 1. J Virol 68:4075–4079.

36. Stein BS, Engleman EG. 1990. Intracellular processing of the gp160 HIV-1 envelope precursor. Endoproteolytic cleavage occurs in a cis or medial compartment of the Golgi complex. J Biol Chem 265:2640–2649.

37. Pal R, Hoke GM, Sarngadharan MG. 1989. Role of oligosaccharides in the processing and maturation of envelope glycoproteins of human immunodeficiency virus type 1. Proc Natl Acad Sci U S A 86:3384–3388.

38. Doores KJ, Bonomelli C, Harvey DJ, Vasiljevic S, Dwek RA, Burton DR, Crispin M, Scanlan CN. 2010. Envelope glycans of immunodeficiency virions are almost entirely oligomannose antigens. Proc Natl Acad Sci U S A 107:13800–13805.

39. Go EP, Ding H, Zhang S, Ringe RP, Nicely N, Hua D, Steinbock RT, Golabek M, Alin J, Alam SM, Cupo A, Haynes BF, Kappes JC, Moore JP, Sodroski JG, Desaire H. 2017. Glycosylation benchmark profile for HIV-1 envelope glycoprotein production based on eleven Env trimers. J Virol 91:e02428–16.

40. Go EP, Herschhorn A, Gu C, Castillo-Menendez L, Zhang S, Mao Y, Chen H, Ding H, Wakefield JK, Hua D, Liao HX, Kappes JC, Sodroski J, Desaire H. 2015. Comparative analysis of the glycosylation profiles of membrane-anchored HIV-1 envelope glycoprotein trimers and soluble gp140. J Virol 89:8245–8257.

41. Geyer H, Holschbach C, Hunsmann G, Schneider J. 1988. Carbohydrates of human immunodeficiency virus. Structures of oligosaccharides linked to the envelope glycoprotein 120. J Biol Chem 263:11760–11767.

42. Munro JB, Gorman J, Ma X, Zhou Z, Arthos J, Burton DR, Koff WC, Courter JR, Smith AB, 3rd, Kwong PD, Blanchard SC, Mothes W. 2014. Conformational dynamics of single HIV-1 envelope trimers on the surface of native virions. Science 346:759–763.

43. Fouts TR, Binley JM, Trkola A, Robinson JE, Moore JP. 1997. Neutralization of the human immunodeficiency virus type 1 primary isolate JR-FL by human monoclonal antibodies correlates with antibody binding to the oligomeric form of the envelope glycoprotein complex. J Virol 71:2779–2785.

44. York J, Follis KE, Trahey M, Nyambi PN, Zolla-Pazner S, Nunberg JH. 2001. Antibody binding and neutralization of primary and T-cell line-adapted isolates of human immunodeficiency virus type 1. J Virol 75:2741–2752.

45. Haim H, Salas I, McGee K, Eichelberger N, Winter E, Pacheco B, Sodroski J. 2013. Modeling virus- and antibody-specific factors to predict human immunodeficiency virus neutralization efficiency. Cell Host Microbe 14:547–558.

46. Guttman M, Cupo A, Julien JP, Sanders RW, Wilson IA, Moore JP, Lee KK. 2015. Antibody potency relates to the ability to recognize the closed, pre-fusion form of HIV Env. Nat Commun 6:6144.

47. Kwong PD, Doyle ML, Casper DJ, Cicala C, Leavitt SA, Majeed S, Steenbeke TD, Venturi M, Chaiken I, Fung M, Katinger H, Parren PW, Robinson J, Van Ryk D, Wang L, Burton DR, Freire E, Wyatt R, Sodroski J, Hendrickson WA, Arthos J. 2002. HIV-1 evades antibody-mediated neutralization through conformational masking of receptor-binding sites. Nature 420:678–682.

48. Herschhorn A, Ma X, Gu C, Ventura JD, Castillo-Menendez L, Melillo B, Terry DS, Smith AB, 3rd, Blanchard SC, Munro JB, Mothes W, Finzi A, Sodroski J. 2016. Release of gp120 restraints leads to an entry-competent intermediate state of the HIV-1 envelope glycoproteins. MBio 7:e01598–16.

49. Ma X, Terry DS, Gorman J, Hong X, Zhou Z, Zhao H, Altman RB, Arthos J, Kwong PD, Blanchard SC, Mothes W, Munro JB. 2018. HIV-1 Env trimer opens through an asymmetric intermediate in which individual protomers adopt distinct conformations. eLife 7:e34271.

50. Furuta RA, Wild CT, Weng Y, Weiss CD. 1998. Capture of an early fusion-active conformation of HIV-1 gp41. Nat Struct Biol 5:276–279.

51. Koshiba T, Chan DC. 2003. The prefusogenic intermediate of HIV-1 gp41 contains exposed C-peptide regions. J Biol Chem 278:7573–7579.

52. He Y, Vassell R, Zaitseva M, Nguyen N, Yang Z, Weng Y, Weiss CD. 2003. Peptides trap the human immunodeficiency virus type 1 envelope glycoprotein fusion intermediate at two sites. J Virol 77:1666–1671.

53. Si Z, Madani N, Cox JM, Chruma JJ, Klein JC, Schon A, Phan N, Wang L, Biorn AC, Cocklin S, Chaiken I, Freire E, Smith AB, 3rd, Sodroski JG. 2004. Small-molecule inhibitors of HIV-1 entry block receptor-induced conformational changes in the viral envelope glycoproteins. Proc Natl Acad Sci U S A 101:5036–5041.

54. Herschhorn A, Gu C, Moraca F, Ma X, Farrell M, Smith AB, 3rd, Pancera M, Kwong PD, Schon A, Freire E, Abrams C, Blanchard SC, Mothes W, Sodroski JG. 2017. The beta20-beta21 of gp120 is a regulatory switch for HIV-1 Env conformational transitions. Nat Commun 8:1049.

55. Castillo-Menendez LR, Nguyen HT, Sodroski J. 2019. Conformational differences between functional human immunodeficiency virus (HIV-1) envelope glycoprotein trimers and stabilized soluble trimers. J Virol 93:e01709–18.

56. Trkola A, Dragic T, Arthos J, Binley JM, Olson WC, Allaway GP, Cheng-Mayer C, Robinson J, Maddon PJ, Moore JP. 1996. CD4-dependent, antibody-sensitive interactions between HIV-1 and its co-receptor CCR-5. Nature 384:184–187.

57. Wu L, Gerard NP, Wyatt R, Choe H, Parolin C, Ruffing N, Borsetti A, Cardoso AA, Desjardin E, Newman W, Gerard C, Sodroski J. 1996. CD4-induced interaction of primary HIV-1 gp120 glycoproteins with the chemokine receptor CCR-5. Nature 384:179–83.

58. Berger EA, Murphy PM, Farber JM. 1999. Chemokine receptors as HIV-1 coreceptors: roles in viral entry, tropism, and disease. Annu Rev Immunol 17:657–700.

59. Ivan B, Sun Z, Subbaraman H, Friedrich N, Trkola A. 2019. CD4 occupancy triggers sequential pre-fusion conformational states of the HIV-1 envelope trimer with relevance for broadly neutralizing antibody activity. PLoS Biol 17:e3000114.

60. Kuhmann SE, Platt EJ, Kozak SL, Kabat D. 2000. Cooperation of multiple CCR5 coreceptors is required for infections by human immunodeficiency virus type 1. J Virol 74:7005–7015.

61. Melikyan GB, Markosyan RM, Hemmati H, Delmedico MK, Lambert DM, Cohen FS. 2000. Evidence that the transition of HIV-1 gp41 into a six-helix bundle, not the bundle configuration, induces membrane fusion. J Cell Biol 151:413–423.

62. Wilen CB, Tilton JC, Doms RW. 2012. Molecular mechanisms of HIV entry. In: Rossmann M, Rao V (eds.), Viral Molecular Machines. Advances in Experimental Medicine and Biology, vol 726. Springer, Boston, MA.

63. Kwong PD, Wyatt R, Robinson J, Sweet RW, Sodroski J, Hendrickson WA. 1998. Structure of an HIV gp120 envelope glycoprotein in complex with the CD4 receptor and a neutralizing human antibody. Nature 393:648–659.

64. Wyatt R, Kwong PD, Desjardins E, Sweet RW, Robinson J, Hendrickson WA, Sodroski JG. 1998. The antigenic structure of the HIV gp120 envelope glycoprotein. Nature 393:705–711.

65. Stewart-Jones GB, Soto C, Lemmin T, Chuang GY, Druz A, Kong R, Thomas PV, Wagh K, Zhou T, Behrens AJ, Bylund T, Choi CW, Davison JR, Georgiev IS, Joyce MG, Kwon YD, Pancera M, Taft J, Yang Y, Zhang B, Shivatare SS, Shivatare VS, Lee CC, Wu CY, Bewley CA, Burton DR, Koff WC, Connors M, Crispin M, Baxa U, Korber BT, Wong CH, Mascola JR, Kwong PD. 2016. Trimeric HIV-1-Env structures define glycan shields from Clades A, B, and G. Cell 165:813–826.

66. Kuiken C, Foley B, Marx P, Wolinsky S, Leitner T, Hahn B, McCutchan F, Korber B. HIV Sequence Compendium 2013. Los Alamos HIV Sequence Database.

67. Wei X, Decker JM, Wang S, Hui H, Kappes JC, Wu X, Salazar-Gonzalez JF, Salazar MG, Kilby JM, Saag MS, Komarova NL, Nowak MA, Hahn BH, Kwong PD, Shaw GM. 2003. Antibody neutralization and escape by HIV-1. Nature 422:307–312.

68. Decker JM, Bibollet-Ruche F, Wei X, Wang S, Levy DN, Wang W, Delaporte E, Peeters M, Derdeyn CA, Allen S, Hunter E, Saag MS, Hoxie JA, Hahn BH, Kwong PD, Robinson JE, Shaw GM. 2005. Antigenic conservation and immunogenicity of the HIV coreceptor binding site. J Exp Med 201:1407–1419.

69. Alsahafi N, Bakouche N, Kazemi M, Richard J, Ding S, Bhattacharyya S, Das D, Anand SP, Prevost J, Tolbert WD, Lu H, Medjahed H, Gendron-Lepage G, Ortega Delgado GG, Kirk S, Melillo B, Mothes W, Sodroski J, Smith AB, 3rd, Kaufmann DE, Wu X, Pazgier M, Rouiller I, Finzi A, Munro JB. 2019. An asymmetric opening of HIV-1 envelope mediates antibody-dependent cellular cytotoxicity. Cell Host Microbe 25:578–587 e5.

70. Labrijn AF, Poignard P, Raja A, Zwick MB, Delgado K, Franti M, Binley J, Vivona V, Grundner C, Huang CC, Venturi M, Petropoulos CJ, Wrin T, Dimitrov DS, Robinson J, Kwong PD, Wyatt RT, Sodroski J, Burton DR. 2003. Access of antibody molecules to the conserved coreceptor binding site on glycoprotein gp120 is sterically restricted on primary human immunodeficiency virus type 1. J Virol 77:10557–10565.

71. Moore PL, Ranchobe N, Lambson BE, Gray ES, Cave E, Abrahams MR, Bandawe G, Mlisana K, Abdool Karim SS, Williamson C, Morris L, CAPRISA 002 study, NIAID Center for HIV/AIDS Vaccine Immunology (CHAVI). 2009. Limited neutralizing antibody specificities drive neutralization escape in early HIV-1 subtype C infection. PLoS Pathog 5:e1000598.

72. Zhang S, Nguyen HT, Ding H, Wang J, Zou S, Liu L, Guha D, Gabuzda D, Ho D, Kappes JC, Sodroski J. Dual pathways of human immunodeficiency virus (HIV-1) envelope glycoprotein trafficking modulate the selective exclusion of uncleaved oligomers from virions. J Virol 95(3):e01369–20.

73. Herrera C, Klasse PJ, Michael E, Kake S, Barnes K, Kibler CW, Campbell-Gardener L, Si Z, Sodroski J, Moore JP, Beddows S. 2005. The impact of envelope glycoprotein cleavage on the antigenicity, infectivity, and neutralization sensitivity of Env-pseudotyped human immunodeficiency virus type 1 particles. Virology 338:154–172.

74. Pancera M, Wyatt R. 2005. Selective recognition of oligomeric HIV-1 primary isolate envelope glycoproteins by potently neutralizing ligands requires efficient precursor cleavage. Virology 332:145–156.

75. Chakrabarti BK, Pancera M, Phogat S, O’Dell S, McKee K, Guenaga J, Robinson J, Mascola J, Wyatt RT. 2011. HIV type 1 Env precursor cleavage state affects recognition by both neutralizing and nonneutralizing gp41 antibodies. AIDS Res Hum Retroviruses 27:877–887.

76. Chakrabarti BK, Walker LM, Guenaga JF, Ghobbeh A, Poignard P, Burton DR, Wyatt RT. 2011. Direct antibody access to the HIV-1 membrane-proximal external region positively correlates with neutralization sensitivity. J Virol 85:8217–8226.

77. Li Y, O’Dell S, Wilson R, Wu X, Schmidt SD, Hogerkorp CM, Louder MK, Longo NS, Poulsen C, Guenaga J, Chakrabarti BK, Doria-Rose N, Roederer M, Connors M, Mascola JR, Wyatt RT. 2012. HIV-1 neutralizing antibodies display dual recognition of the primary and coreceptor binding sites and preferential binding to fully cleaved envelope glycoproteins. J Virol 86:11231–11241.

78. Castillo-Menendez LR, Witt K, Espy N, Princiotto A, Madani N, Pacheco B, Finzi A, Sodroski J. 2018. Comparison of uncleaved and mature human immunodeficiency virus membrane envelope glycoprotein trimers. J Virol 92:e00277–18.

79. Zou S, Zhang S, Gaffney A, Ding H, Lu M, Grover JR, Farrell M, Nguyen HT, Zhao C, Anang S, Zhao M, Mohammadi M, Blanchard SC, Abrams C, Madani N, Mothes W, Kappes JC, Smith AB, 3rd, Sodroski J. 2020. Long-acting BMS-378806 analogues stabilize the State-1 conformation of the human immunodeficiency virus type 1 envelope glycoproteins. J Virol 94:e00148–20.

80. Haim H, Salas I, Sodroski J. 2013. Proteolytic processing of the human immunodeficiency virus envelope glycoprotein precursor decreases conformational flexibility. J Virol 87:1884–1889.

81. Lu M, Ma X, Reichard N, Terry DS, Arthos J, Smith AB, 3rd, Sodroski JG, Blanchard SC, Mothes W. 2020. Shedding-resistant HIV-1 envelope glycoproteins adopt downstream conformations that remain responsive to conformation-preferring ligands. J Virol 94:e00597–20.

82. Wang Q, Finzi A, Sodroski J. 2020. The Conformational States of the HIV-1 Envelope Glycoproteins. Trends Microbiol 28:655–667.

83. Wibmer CK, Bhiman JN, Gray ES, Tumba N, Abdool Karim SS, Williamson C, Morris L, Moore PL. 2013. Viral escape from HIV-1 neutralizing antibodies drives increased plasma neutralization breadth through sequential recognition of multiple epitopes and immunotypes. PLoS Pathog 9:e1003738.

84. Gray ES, Taylor N, Wycuff D, Moore PL, Tomaras GD, Wibmer CK, Puren A, DeCamp A, Gilbert PB, Wood B, Montefiori DC, Binley JM, Shaw GM, Haynes BF, Mascola JR, Morris L. 2009. Antibody specificities associated with neutralization breadth in plasma from human immunodeficiency virus type 1 subtype C-infected blood donors. J Virol 83:8925–8937.

85. Sather DN, Armann J, Ching LK, Mavrantoni A, Sellhorn G, Caldwell Z, Yu X, Wood B, Self S, Kalams S, Stamatatos L. 2009. Factors associated with the development of cross-reactive neutralizing antibodies during human immunodeficiency virus type 1 infection. J Virol 83:757–769.

86. Klein F, Diskin R, Scheid JF, Gaebler C, Mouquet H, Georgiev IS, Pancera M, Zhou T, Incesu RB, Fu BZ, Gnanapragasam PN, Oliveira TY, Seaman MS, Kwong PD, Bjorkman PJ, Nussenzweig MC. 2013. Somatic mutations of the immunoglobulin framework are generally required for broad and potent HIV-1 neutralization. Cell 153:126–138.

87. Walker LM, Simek MD, Priddy F, Gach JS, Wagner D, Zwick MB, Phogat SK, Poignard P, Burton DR. 2010. A limited number of antibody specificities mediate broad and potent serum neutralization in selected HIV-1 infected individuals. PLoS Pathog 6:e1001028.

88. Gray ES, Madiga MC, Hermanus T, Moore PL, Wibmer CK, Tumba NL, Werner L, Mlisana K, Sibeko S, Williamson C, Abdool Karim SS, Morris L, Team CS. 2011. The neutralization breadth of HIV-1 develops incrementally over four years and is associated with CD4+ T cell decline and high viral load during acute infection. J Virol 85:4828–4840.

89. Corti D, Langedijk JP, Hinz A, Seaman MS, Vanzetta F, Fernandez-Rodriguez BM, Silacci C, Pinna D, Jarrossay D, Balla-Jhagjhoorsingh S, Willems B, Zekveld MJ, Dreja H, O’Sullivan E, Pade C, Orkin C, Jeffs SA, Montefiori DC, Davis D, Weissenhorn W, McKnight A, Heeney JL, Sallusto F, Sattentau QJ, Weiss RA, Lanzavecchia A. 2010. Analysis of memory B cell responses and isolation of novel monoclonal antibodies with neutralizing breadth from HIV-1-infected individuals. PLoS One 5:e8805.

90. Wu X, Zhou T, Zhu J, Zhang B, Georgiev I, Wang C, Chen X, Longo NS, Louder M, McKee K, O’Dell S, Perfetto S, Schmidt SD, Shi W, Wu L, Yang Y, Yang ZY, Yang Z, Zhang Z, Bonsignori M, Crump JA, Kapiga SH, Sam NE, Haynes BF, Simek M, Burton DR, Koff WC, Doria-Rose NA, Connors M, Program NCS, Mullikin JC, Nabel GJ, Roederer M, Shapiro L, Kwong PD, Mascola JR. 2011. Focused evolution of HIV-1 neutralizing antibodies revealed by structures and deep sequencing. Science 333:1593–1602.

91. Hraber P, Seaman MS, Bailer RT, Mascola JR, Montefiori DC, Korber BT. 2014. Prevalence of broadly neutralizing antibody responses during chronic HIV-1 infection. AIDS 28:163–169.

92. Upadhyay C, Mayr LM, Zhang J, Kumar R, Gorny MK, Nadas A, Zolla-Pazner S, Hioe CE. 2014. Distinct mechanisms regulate exposure of neutralizing epitopes in the V2 and V3 loops of HIV-1 envelope. J Virol 88:12853–12865.

93. Zolla-Pazner S, Cohen SS, Boyd D, Kong XP, Seaman M, Nussenzweig M, Klein F, Overbaugh J, Totrov M. 2016. Structure/function studies involving the V3 region of the HIV-1 envelope delineate multiple factors that affect neutralization sensitivity. J Virol 90:636–649.

94. Powell RLR, Totrov M, Itri V, Liu X, Fox A, Zolla-Pazner S. 2017. Plasticity and epitope exposure of the HIV-1 envelope trimer. J Virol 91:e00410–17.

95. Zhang S, Wang WL, Chen S, Luy M, Go EP, Steinbock RT, Ding H, Desaire H, Kappes JC, Sodroski J, Mao Y. 2018. Structural insights into the conformational plasticity of the full-length trimeric HIV-1 envelope glycoprotein precursor. bioRxiv doi: 10.1101/288472.

96. Scheres SH. 2012. RELION: Implementation of a Bayesian approach to cryo-EM structure determination. J Struct Biol 180:519–30.

97. Julien JP, Cupo A, Sok D, Stanfield RL, Lyumkis D, Deller MC, Klasse PJ, Burton DR, Sanders RW, Moore JP, Ward AB, Wilson IA. 2013. Crystal structure of a soluble cleaved HIV-1 envelope trimer. Science 342:1477–1483.

98. Lyumkis D, Julien JP, de Val N, Cupo A, Potter CS, Klasse PJ, Burton DR, Sanders RW, Moore JP, Carragher B, Wilson IA, Ward AB. 2013. Cryo-EM structure of a fully glycosylated soluble cleaved HIV-1 envelope trimer. Science 342:1484–1490.

99. Pancera M, Zhou T, Druz A, Georgiev IS, Soto C, Gorman J, Huang J, Acharya P, Chuang GY, Ofek G, Stewart-Jones GB, Stuckey J, Bailer RT, Joyce MG, Louder MK, Tumba N, Yang Y, Zhang B, Cohen MS, Haynes BF, Mascola JR, Morris L, Munro JB, Blanchard SC, Mothes W, Connors M, Kwong PD. 2014. Structure and immune recognition of trimeric pre-fusion HIV-1 Env. Nature 514:455–461.

100. Bartesaghi A, Merk A, Borgnia MJ, Milne JL, Subramaniam S. 2013. Prefusion structure of trimeric HIV-1 envelope glycoprotein determined by cryo-electron microscopy. Nat Struct Mol Biol 20:1352–1357.

101. Guenaga J, de Val N, Tran K, Feng Y, Satchwell K, Ward AB, Wyatt RT. 2015. Well-ordered trimeric HIV-1 subtype B and C soluble spike mimetics generated by negative selection display native-like properties. PLoS Pathog 11:e1004570.

102. Guenaga J, Dubrovskaya V, Val N, Sharma SK, Carrette B, Ward AB, Wyatt RT. 2015. Structure-guided redesign increases the propensity of HIV Env to generate highly stable soluble trimers. J Virol 90:2806–2817.

103. Georgiev IS, Joyce MG, Yang Y, Sastry M, Zhang B, Baxa U, Chen RE, Druz A, Lees CR, Narpala S, Schön A, Van Galen J, Chuang GY, Gorman J, Harned A, Pancera M, Stewart-Jones GB, Cheng C, Freire E, McDermott AB, Mascola JR, Kwong PD. 2015. Single-chain soluble BG505.SOSIP gp140 trimers as structural and antigenic mimics of mature closed HIV-1 Env. J Virol 89:5318–5329.

104. Kwon YD, Pancera M, Acharya P, Georgiev IS, Crooks ET, Gorman J, Joyce MG, Guttman M, Ma X, Narpala S, Soto C, Terry DS, Yang Y, Zhou T, Ahlsen G, Bailer RT, Chambers M, Chuang GY, Doria-Rose NA, Druz A, Hallen MA, Harned A, Kirys T, Louder MK, O’Dell S, Ofek G, Osawa K, Prabhakaran M, Sastry M, Stewart-Jones GB, Stuckey J, Thomas PV, Tittley T, Williams C, Zhang B, Zhao H, Zhou Z, Donald BR, Lee LK, Zolla-Pazner S, Baxa U, Schon A, Freire E, Shapiro L, Lee KK, Arthos J, Munro JB, Blanchard SC, Mothes W, Binley JM, McDermott AB, Mascola JR, Kwong PD. 2015. Crystal structure, conformational fixation and entry-related interactions of mature ligand-free HIV-1 Env. Nat Struct Mol Biol 22:522–531.

105. Lee JH, Ozorowski G, Ward AB. 2016. Cryo-EM structure of a native, fully glycosylated, cleaved HIV-1 envelope trimer. Science 351:1043–1048.

106. Gristick HB, von Boehmer L, West AP, Jr., Schamber M, Gazumyan A, Golijanin J, Seaman MS, Fatkenheuer G, Klein F, Nussenzweig MC, Bjorkman PJ. 2016. Natively glycosylated HIV-1 Env structure reveals new mode for antibody recognition of the CD4-binding site. Nat Struct Mol Biol 23:906–915.

107. Pan J, Peng H, Chen B, Harrison SC. 2020. Cryo-EM structure of full-length HIV-1 Env bound with the Fab of antibody PG16. J Mol Biol 432:1158–1168.

108. Torrents de la Pena A, Rantalainen K, Cottrell CA, Allen JD, van Gils MJ, Torres JL, Crispin M, Sanders RW, Ward AB. 2019. Similarities and differences between native HIV-1 envelope glycoprotein trimers and stabilized soluble trimer mimetics. PLoS Pathog 15:e1007920.

109. Pacheco B, Alsahafi N, Debbeche O, Prévost J, Ding S, Chapleau JP, Herschhorn A, Madani N, Princiotto A, Melillo B, Gu C, Zeng X, Mao Y, Smith AB 3rd, Sodroski J, Finzi A. 2017. Residues in the gp41 ectodomain regulate HIV-1 envelope glycoprotein conformational transitions induced by gp120-directed inhibitors. J Virol 91:e02219–16.

110. Sen J, Yan T, Wang J, Rong L, Tao L, Caffrey M. 2010. Alanine scanning mutagenesis of HIV-1 gp41 heptad repeat 1: insight into the gp120-gp41 interaction. Biochemistry 49:5057–5065.

111. Keller PW, Morrison O, Vassell R, Weiss CD. 2018. HIV-1 gp41 residues modulate CD4-induced conformational changes in the envelope glycoprotein and evolution of a relaxed conformation of gp120. J Virol 92:e00583–18.

112. Dey AK, David KB, Klasse PJ, Moore JP. 2007. Specific amino acids in the N-terminus of the gp41 ectodomain contribute to the stabilization of a soluble, cleaved gp140 envelope glycoprotein from human immunodeficiency virus type 1. Virology 360:199–208.

113. Ringe RP, Sanders RW, Yasmeen A, Kim HJ, Lee JH, Cupo A, Korzun J, Derking R, van Montfort T, Julien JP, Wilson IA, Klasse PJ, Ward AB, Moore JP. 2013. Cleavage strongly influences whether soluble HIV-1 envelope glycoprotein trimers adopt a native-like conformation. Proc Natl Acad Sci U S A. 110:18256–18261.

114. Ringe RP, Yasmeen A, Ozorowski G, Go EP, Pritchard LK, Guttman M, Ketas TA, Cottrell CA, Wilson IA, Sanders RW, Cupo A, Crispin M, Lee KK, Desaire H, Ward AB, Klasse PJ, Moore JP. 2015. Influences on the design and purification of soluble, recombinant native-like HIV-1 envelope glycoprotein trimers. J Virol 89:12189–12210.

115. Ringe RP, Colin P, Torres JL, Yasmeen A, Lee WH, Cupo A, Ward AB, Klasse PJ, Moore JP. 2019. SOS and IP modifications predominantly affect the yield but not other properties of SOSIP.664 HIV-1 Env glycoprotein trimers. J Virol 94:e01521–19.

116. Yang Z, Wang H, Liu AZ, Gristick HB, Bjorkman PJ. 2019. Asymmetric opening of HIV-1 Env bound to CD4 and a coreceptor-mimicking antibody. Nat Struct Mol Biol 26:1167–1175. Erratum in: Nat Struct Mol Biol 2020 27:105.

117. Pritchard LK, Vasiljevic S, Ozorowski G, Seabright GE, Cupo A, Ringe R, Kim HJ, Sanders RW, Doores KJ, Burton DR, Wilson IA, Ward AB, Moore JP, Crispin M. 2015. Structural constraints determine the glycosylation of HIV-1 envelope trimers. Cell Rep 11:1604–1613.

118. Pancera M, Lai YT, Bylund T, Druz A, Narpala S, O’Dell S, Schön A, Bailer RT, Chuang GY, Geng H, Louder MK, Rawi R, Soumana DI, Finzi A, Herschhorn A, Madani N, Sodroski J, Freire E, Langley DR, Mascola JR, McDermott AB, Kwong PD. 2017. Crystal structures of trimeric HIV envelope with entry inhibitors BMS-378806 and BMS-626529. Nat Chem Biol 13:1115–1122.

119. Finzi A, Xiang SH, Pacheco B, Wang L, Haight J, Kassa A, Danek B, Pancera M, Kwong PD, Sodroski J. 2010. Topological layers in the HIV-1 gp120 inner domain regulate gp41 interaction and CD4-triggered conformational transitions. Mol Cell 37:656–667.

120. Chen J, Lee KH, Steinhauer DA, Stevens DJ, Skehel JJ, Wiley DC. 1998. Structure of the hemagglutinin precursor cleavage site, a determinant of influenza pathogenicity and the origin of the labile conformation. Cell 95:409–417.

121. Lu M, Ma X, Castillo-Menendez LR, Gorman J, Alsahafi N, Ermel U, Terry DS, Chambers M, Peng D, Zhang B, Zhou T, Reichard N, Wang K, Grover J, Carman BP, Gardner MR, Nikíc-Spiegel I, Sugawara A, Arthos J, Lemke EA, Smith AB, 3rd, Farzan M, Abrams C, Munro JB, McDermott AB, Finzi A, Kwong PD, Blanchard SC, Sodroski JG, Mothes, W. 2019. Associating HIV-1 envelope glycoprotein structures with states on virus observed by smFRET. Nature 568:415–419.

122. Salimi H, Johnson J, Flores MG, Zhang MS, O’Malley Y, Houtman JC, Schlievert PM, Haim H. 2020. The lipid membrane of HIV-1 stabilizes the viral envelope glycoproteins and modulates their sensitivity to antibody neutralization. J Biol Chem 295:348–362.

123. Mao Y, Wang L, Gu C, Herschhorn A, Xiang SH, Haim H, Yang X, Sodroski J. 2012. Subunit organization of the membrane-bound HIV-1 envelope glycoprotein trimer. Nat Struct Mol Biol 19:893–899.

124. Mao Y, Wang L, Gu C, Herschhorn A, Désormeaux A, Finzi A, Xiang SH, Sodroski JG. 2013. Molecular architecture of the uncleaved HIV-1 envelope glycoprotein trimer. Proc Natl Acad Sci U S A 110:12438–12443.

125. Wu Z, Zhang S, Wang WL, Ma Y, Dong Y, Mao Y. 2020. Deep manifold learning reveals hidden dynamics of proteasome autoregulation. bioRxiv doi: 10.1101/2020.12.22.423932.

126. Punjani A, Fleet DJ. 2020. 3D variability analysis: Resolving continuous flexibility and discrete heterogeneity from single particle cryo-EM. bioRxiv doi: 10.1101/2020.04.08.032466.

127. McCune JM, Rabin LB, Feinberg MB, Lieberman M, Kosek JC, Reyes GR, Weissman IL. 1988. Endoproteolytic cleavage of gp160 is required for the activation of human immunodeficiency virus. Cell 53:55–67.

128. Moyer MP, Huot RI, Ramirez A Jr, Joe S, Meltzer MS, Gendelman HE. 1990. Infection of human gastrointestinal cells by HIV-1. AIDS Res Hum Retroviruses 6:1409–1415.

129. Juette MF, Terry DS, Wasserman MR, Altman RB, Zhou Z, Zhao H, Blanchard SC. 2016. Single-molecule imaging of non-equilibrium molecular ensembles on the millisecond timescale. Nat Methods 13:341–344.

130. Potter CS, Chu H, Frey B, Green C, Kisseberth N, Madden TJ, Miller KL, Nahrstedt K, Pulokas J, Reilein A, Tcheng D, Weber D, Carragher B. 1999. Leginon: a system for fully automated acquisition of 1000 electron micrographs a day. Ultramicroscopy 77:153–161.

131. Cheng A, Negro C, Bruhn JF, Rice WJ, Dallakyan S, Eng ET, Waterman DG, Potter CS, Carragher B. 2021. Leginon: New features and applications. Protein Sci 30:136–150.

132. Mastronarde DN. 2005. Automated electron microscope tomography using robust prediction of specimen movements. J Struct Biol 152:36–51.

133. Zheng SQ, Palovcak E, Armache JP, Verba KA, Cheng Y, Agard DA. 2017. MotionCor2: anisotropic correction of beam-induced motion for improved cryo-electron microscopy. Nat Methods 14:331–332.

134. Rohou A, Grigorieff N. 2015. CTFFIND4: Fast and accurate defocus estimation from electron micrographs. J Struct Biol 192:216–221.

135. Zhang K. Gctf: 2016. Real-time CTF determination and correction. J Struct Biol 193:1–12.

136. Zhu Y, Ouyang Q, Mao Y. 2017. A deep convolutional neural network approach to single-particle recognition in cryo-electron microscopy. BMC Bioinformatics 18:348.

137. Zivanov J, Nakane T, Forsberg BO, Kimanius D, Hagen WJ, Lindahl E, Scheres SH. 2018. New tools for automated high-resolution cryo-EM structure determination in RELION-3. Elife 7:e42166.

138. Wu J, Ma YB, Congdon C, Brett B, Chen S, Xu Y, Ouyang Q, Mao Y. 2017. Massively parallel unsupervised single-particle cryo-EM data clustering via statistical manifold learning. PLoS One 12:e0182130.

139. Pettersen, E. F. et al. 2004. UCSF Chimera—a visualization system for exploratory research and analysis. J Comput Chem 25:1605–1612.

140. Casañal A, Lohkamp B, Emsley P. 2020. Current developments in Coot for macromolecular model building of electron cryo-microscopy and crystallographic data. Protein Sci 29:1069–1078.

141. Afonine PV, Poon BK, Read RJ, Sobolev OV, Terwilliger TC, Urzhumtsev A, Adams PD. 2018. Real-space refinement in PHENIX for cryo-EM and crystallography. Acta Crystallogr D Struct Biol 74:531–544.

142. The PyMOL Molecular Graphics System Version 1.8 (Schrödinger, LLC).

143. Kucukelbir A, Sigworth FJ, Tagare HD. 2014. Quantifying the local resolution of cryo-EM density maps. Nat Methods 11:63–65.

